# Amygdala beta bursts modulate hippocampal neuronal dynamics during human emotional processing

**DOI:** 10.1101/2025.09.26.678792

**Authors:** Yajun Zhou, Yuqing Huang, Giana F. Pittaro, Behzad Zareian, Ayman Aljishi, Alondra P. Toro, Zion Zibly, Sami Obaid, Adithya Sivaraju, Ilan Harpaz-Rotem, Alfred Kaye, John Krystal, Kevin Sheth, Xiaosi Gu, Christopher Pittenger, Eyiyemisi C. Damisah

## Abstract

The amygdala and hippocampus are key structures in emotional processing, yet the transient neural dynamics coordinating their interaction remain poorly understood. We simultaneously recorded single-neuron activity and local field potentials from these regions in epilepsy patients performing an emotional image-rating task. Neurons in both areas exhibited firing rate changes predictive of subjective emotional valence. While conventional spectral analysis revealed no valence-specific patterns, beta bursts (13–30 Hz)—transient, high-power events-were uniquely modified by task features. In both regions, beta bursts were associated with increased gamma amplitude and enhanced phase coherence, with beta-gamma phase-amplitude coupling capturing emotion-related dynamics. Critically, amygdala beta bursts were associated with strong suppression of hippocampal firing, coinciding with interneuron activation, during negative valence/highest arousal processing, whereas hippocampal bursts showed no reciprocal influence. These findings suggest that beta bursts may represent a temporally precise neural correlate of emotional appraisal and a candidate mechanism for targeted neuromodulation in mood disorders.

## 1 Introduction

Emotion appraisal and regulation are fundamental to human cognition and behavior. The amygdala, a key region for detecting and evaluating emotionally salient stimuli, communicates extensively with the hippocampus, which provides contextual information to support the encoding and modulation of emotional experiences[1–4]. Through reciprocal connections, these structures prioritize emotionally salient information and shape future responses to affective stimuli [1, 5–8]. In particular, the bidirectional circuit between the basolateral amygdala (BLA) and the ventral hippocampus (vHPC) has been implicated in a range of emotion-related tasks, especially in the context of negative emotional behaviors [9–12]. Disruption of the amygdala-hippocampal circuit is observed in various neuropsychiatric conditions, including major depression, post-traumatic stress disorder, and pathological anxiety, which collectively affect millions of individuals worldwide[9, 10, 13–22]. Despite this, the precise temporal dynamics that coordinate rapid amygdala–hippocampus interactions at the single-neuron level in humans during emotional processing remain poorly understood.

Recent studies have highlighted the importance of neural oscillations in modulating neuronal activity [23–25]. In particular, beta oscillations (13–30 Hz) have been implicated in mediating long-range, state-dependent signaling for movement control, sensorimotor integration, working memory, and decision making [26–29]. However, the role of beta oscillations in emotion processing remains controversial: while some human EEG, MEG studies report decreases in beta power during emotional processing [30–35], others observe increases, especially during negative affect or anxiety [36–38]. These inconsistencies may arise from differences in species, methodologies, and behavioral paradigms. Moreover, conventional spectral analyses that average neural activity over extended time windows may obscure rapid, millisecond-scale dynamics that are critical for emotional processing [39, 40].

Beta oscillations often manifest not as sustained rhythms but as bursts—transient, high-power events that coordinate neural activity with millisecond precision [40–43]. In sensorimotor cortices, both human EEG and non-human primate intracranial studies have shown that individual beta bursts predict behavior more accurately than averaged beta power [44–46].In prefrontal networks, intracranial studies in non-human primates indicate that the timing of bursts shapes information routing across regions [39, 47]. Notably, recent mechanistic work in rodent anxiety models [18] has extended this framework to the amygdala-hippocampus circuit, demonstrating that beta synchrony, driven by somatostatin interneurons, strictly regulates circuit excitability. Within these local microcircuits, such inhibitory interneurons typically function to constrain the activity of excitatory pyramidal cells. These findings suggest that beta bursts may serve a specific ‘inhibitory gating’ function [48–50], distinct from the information transfer and plasticity typically attributed to slower oscillations such as theta and alpha rhythms [51, 52]. Thus, emotion processing may depend on the precise timing of transient events, which could be obscured by analyses that focus solely on broadband oscillatory power.

Motivated by the absence of emotional-state correlates in conventional band-power analyses, we hypothesized that transient beta bursts are associated with coordinated modulation of amygdala-hippocampal activity during human emotional processing. To test this, we simultaneously recorded single-neuron activity and local field potentials from the amygdala and hippocampus in human subjects performing an affective image rating task. This approach enabled us to examine how population-level oscillatory events relate to the timing of neuronal spiking, providing insight into the circuit and neuron-level mechanisms underlying emotional processing in humans. Our results show that beta bursts modulate high gamma (HG) amplitude and single-neuron firing rates in an emotion-dependent manner, with amygdala bursts coinciding with transient reductions in hippocampal spiking activity during negative emotions. These findings suggest that beta bursts provide a temporal precise mechanism for emotion appraisal and may represent a candidate neuronal biomarker and therapeutic target for neuromodulation.

## 2 Results

### 2.1 Experimental design and behavioral responses

To investigate the neural mechanisms underpinning the coding of emotional valence in humans, we recorded LFPs and single-neuron activity from the hippocampus and amygdala of 15 epilepsy patients. The participants’ demographic and clinical characteristics—including age, sex, handedness, anti-seizure medications, epilepsy etiology, psychiatric medications and scores on the Beck Anxiety Inventory (BAI) [53] and Beck Depression Inventory (BDI) [54]—are detailed in the Methods. On average, participants had BAI scores of 15.25 ± 3.42 (mean ± SEM), indicating mild anxiety, and BDI scores of 14.63 ± 3.04, indicating mild depression, based on standard inventory scales. (Fig. 1a, Table S1 and Table S3).

**Fig. 1:**
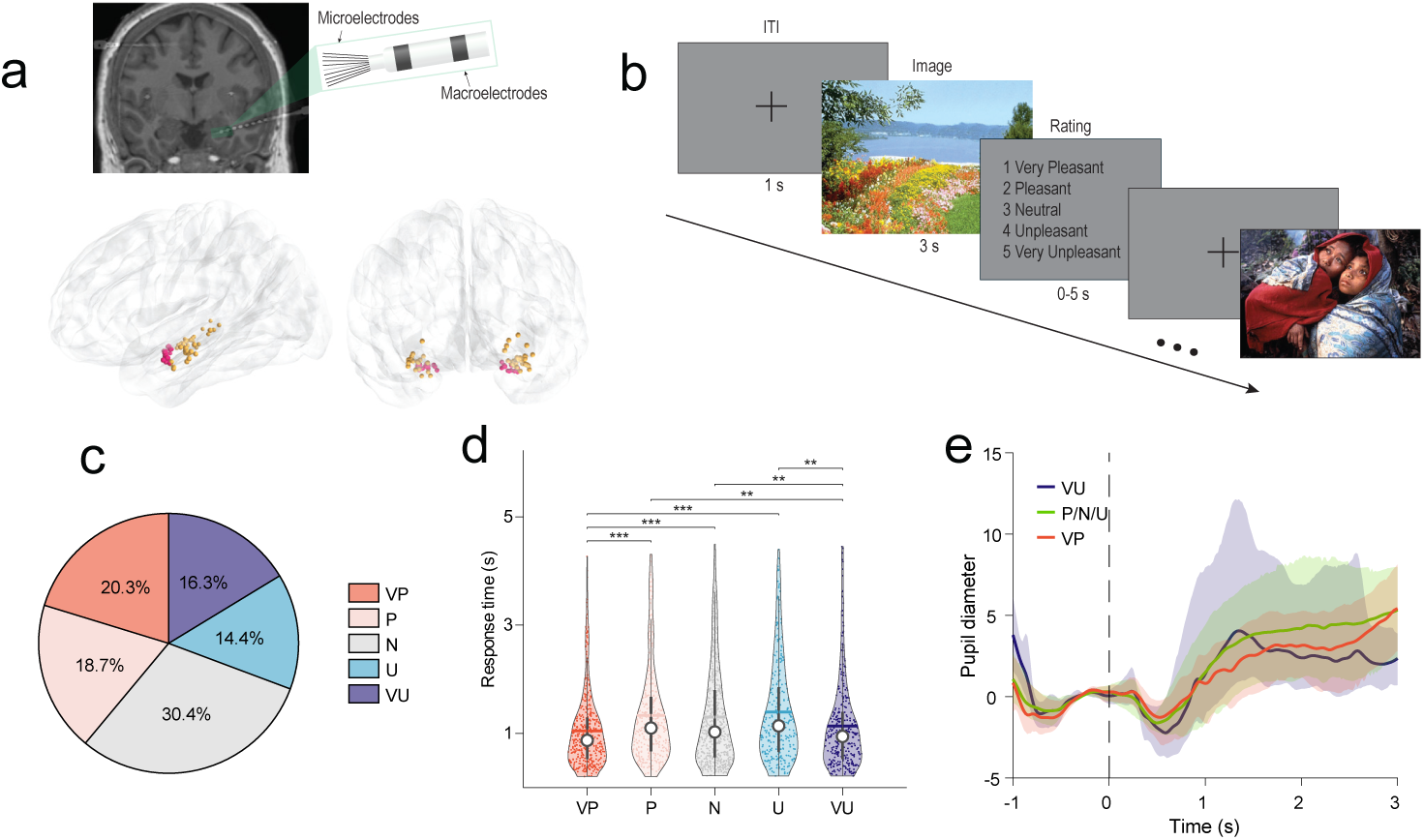
Experimental paradigm and behavioral validation of emotional processing. **a**, Recording setup and electrode localization. Top: Behnke-Fried depth electrode with protruding microwires enabling simultaneous LFP and single-unit recordings. Bottom: Microwire locations across all participants (n=15) shown in MNI space. AMY microwire bundles (n=16, pink), hippocampus (n=23 microwire bundles orange). **b**, Emotional valence rating task. Participants viewed IAPS/NAPS images for 3 s followed by a 5-point valence rating (1=very pleasant, 5=very unpleasant). Inter-trial interval: 1s. **c**, Distribution of valence ratings across all trials (n=2,270). Neutral ratings were most frequent (30.4%), extreme ratings least frequent (very pleasant: 20.3%, very unpleasant: 16.3%). **d**, Response times showed an inverted-U pattern with faster decisions for extreme emotional rating. Violin plots show distribution with median (white dots), IQR (thick bar), and full range (thin lines). ART ANOVA test followed by permutation tests with FDR correction. \*\**p <* 0.01, \*\*\**p <* 0.001. **e**, Pupil diameter responses showed no differences across valence conditions. Shaded areas: ±s.e.m. ART ANOVA: no main effect of condition (*p >* 0.05). AMY: amygdala, HPC: hippocampus. VP: very pleasant, P: pleasant, N: neutral, U: unpleasant, VU: very unpleasant.

Each electrode consisted of macro- and microcontacts, enabling parallel recordings of local field potentials and unit action potentials from the amygdala (n = 29 LFP channels, 82 units) and hippocampus (n = 46 LFP channels, 136 units; Table S4). During the recordings, participants rated images [55][56] on a 5-point valence scale ranging from 1 (very pleasant, VP) to 5 (very unpleasant, VU) following 3-second presentations (Fig. 1b). Across participants, ratings were distributed across all valence categories, with neutral responses being the most common (30.4% ± 12.1%) and extreme ratings less frequent (very pleasant: 20.3% ± 10.0%; very unpleasant: 16.3% ± 10.8%; Fig. 1c). The distribution of ratings was influenced by the semantic category of the images: objects and humans were more often rated as neutral, while animals and scenes were more frequently rated as very pleasant (Fig. S1a). Statistical analysis confirmed significant differences in rating distributions across stimulus categories (aligned ranks transformation (ART) ANOVA, animal: *F* (4, 90) = 6.76*, p* = 8.34 × 10*^−^*^5^, human: *F* (4, 90) = 6.29*, p* = 1.64 × 10*^−^*^4^, scene: *F* (4, 90) = 15.89*, p* = 7.11 × 10*^−^*^10^, object: *F* (4, 95) = 36.62*, p* = 2.22 × 10*^−^*^16^). Permutation test with multiple comparisons correction by FDR revealed that participants rated objects (*p <* 0.01 for 3 vs. all others) and humans (*p <* 0.05 for 3 vs. all others) more frequently as neutral, and rated animals (*p* = 0.005 for 1 vs. 4 and *p <* 0.001 for 1 vs. 5) and scenes (*p <* 0.001 for 1 vs. all others) more frequently as very pleasant.

Response times (RT) revealed a behavioral correlate of emotional processing. We observed an inverted-U pattern (*F*_(4,1958)_ = 10.01, *p* = 5.15 × 10*^−^*^8^), with significantly faster responses for emotional extremes compared to moderate ratings (very pleasant vs. moderate: all *p <* 0.001; very unpleasant vs. moderate: all *p <* 0.01; Fig. 1d). In contrast, we found no significant difference in RT across semantic categories (*F* (3, 1970) = 0.97*, p* = 0.41, Fig S1b). This inverted U pattern suggests faster processing of high-valence stimuli. To determine if these valence effects were confounded by arousal, we analyzed the normative arousal ratings of the images. We found that ‘very unpleasant’ images possessed significantly higher arousal scores compared to neutral or pleasant images in both the IAPS and NAPS datasets (Fig S1d–e, ART ANOVA, *F* (4, 1195) = 546.44*, p* = 2.22 × 10*^−^*^16^ and *F* (4, 1079) = 117.47*, p* = 2.22 × 10*^−^*^16^).

Despite these normative differences, pupil responses recorded from 12 sessions showed a characteristic dilation but did not differ significantly across valence conditions (Fig. 1e, ART ANOVA, *p >* 0.05), or across semantic categories (Fig. S1c, ART ANOVA, *p >* 0.05). We also split trials based on their normative arousal scores (High vs. Low Standardized Arousal). While the “High Arousal” condition elicited a visually larger pupil dilation than the “Low Arousal” condition, this difference did not reach statistical significance (Fig. S1f, ART ANOVA, *p >* 0.05). This suggests that while the ‘very unpleasant’ stimuli are intrinsically more arousing, they did not elicit a differential autonomic pupil response within the short (3 s) window of our task. To further examine whether pupil responses were modulated by arousal, we applied a basis-function analysis that separates early light-reflex and later cognitive components. We observed no significant difference between high- and low-arousal conditions (paired t-test, *p* = 0.173; Bayes Factor *BF*_01_ = 1.44) during the later cognitive component of the pupillary response to standardized arousal ratings. While we do not observe a reliable pupil modulation by arousal, the Bayesian analysis indicates only anecdotal evidence and remains inconclusive with respect to distinguishing between the null and alternative hypotheses. Accordingly, we refer to the “very unpleasant” condition as “very unpleasant/highest arousal” throughout.

### 2.2 Amygdala and hippocampal neurons encode emotional valence with graded selectivity

To investigate how individual neurons in the amygdala and hippocampus process emotional information, we computed trial-wise peristimulus time histograms (PSTH) for each subjective emotional rating. We identified 218 well-isolated neurons (Fig. 2a, Table S4) across amygdala and hippocampus following spike sorting (Table S4). 65.1% (54/82) of neurons in the amygdala and 59.6% (81/136) in the hippocampus exhibited significant modulation of firing rates in response to emotional stimuli (Fig. 2b). Many emotion-responsive neurons in both regions exhibited elevated firing for specific emotional ratings compared to others, with firing rates diverging significantly by 500-800 ms post-stimulus (cluster-based permutation t-test, *p <* 0.05, Fig. 2c-d). While most responsive neurons were selective for single emotional categories, we observed a subset (38.89% in the amygdala and 36.59% in the hippocampus) that responded to adjacent ratings, suggesting a graded rather than categorical representation of valence (Fig. 2b). This partial overlap was more pronounced for moderate emotional ratings than for extreme ratings.

**Fig. 2:**
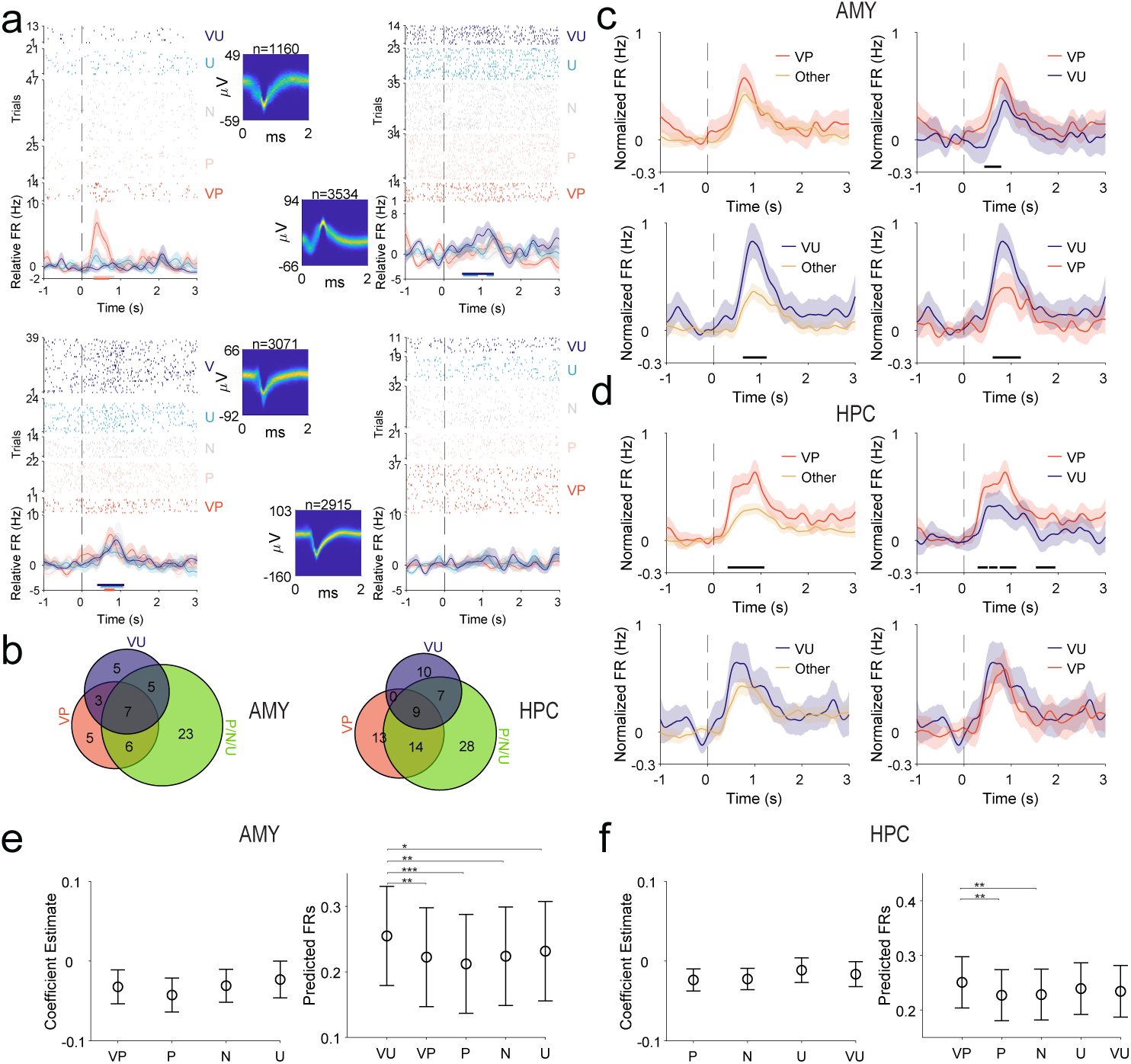
Valence-specific single neuron responses in the human AMY and HPC. **a**, Representative single neurons showing distinct emotional response patterns. Each panel shows raster plots (top) of trials sorted by valence rating and PSTHs (bottom) with baseline-corrected firing rates (± s.e.m.). Four exampler emotional rating associated neuronal responses are shown: Very pleaseant (top left, from HPC), Very unpleasant (top right from AMY), broadly responsive (bottom left, AMY), and non-responsive (bottom right, HPC) neurons. Insets show spike waveforms with amplitude scale bars. **b**, Venn diagrams revealing overlap of valence-responsive neurons in AMY (left, 54 total) and HPC (right, 81 total), with numbers indicating neurons selective for each rating combination. **c**, Population responses of valence-selective neurons in AMY. Top row: very pleasant-responsive neurons (n=13); bottom row: very unpleasant-responsive neurons (n=15). Left panels compare the corresponding trials to all other trials among the responsive neurons; right panels show direct comparison between trials for very pleasant and very unpleasant conditions. Black bars indicate significant differences (cluster-based permutation, *p <* 0.05). **d**, Same analysis for HPC neurons showing very pleasant-responsive (n=27, top) and very unpleasant-responsive (n=12, bottom) populations. **e**, LME results for VU-responsive neurons in AMY. Left: *β* coefficients (mean ± 95% CI) from a population-level LME with valence (VU as reference) as fixed predictors of task-period FR, unit and subject as rondom intercepts. Right: model-predicted FRs (mean ± 95% CI) at the mean baseline FR. **f**, Same as **e**, but for VP-responsive neurons in HPC with VP as the reference. \**p <* 0.05, \*\**p <* 0.01, \*\*\**p <* 0.001.

To evaluate how emotional valence modulated neuronal firing while accounting for subject-level variability, we also analyzed the firing rates of valence-selective neurons using a linear mixed-effects model (LME) with subject and unit as random intercepts, and baseline firing rate and valence as fixed effects. For amygdala neurons responsive to very unpleasant stimuli (Fig. 2e), this analysis revealed a significant overall effect of valence on task-period firing rates (*F* (4, 1617.8) = 5.482, *p* = 0.0022), while baseline firing rate was a strong positive predictor of task-period activity (*β* = −0.077, 95% CI [−0.131, −0.022], *p* = 0.0058). Because baseline and task-period firing rates are measured on the same scale, this coefficient reflects shared firing-rate variability rather than the magnitude of stimulus-evoked responses: neurons with higher spontaneous firing rates tended to exhibit correspondingly higher firing during the task period. Post-hoc analysis showed significantly increased firing for very unpleasant condition compared to very pleasant (*β* = −0.033, 95% CI [−0.054, −0.011], *p* = 0.004), pleasant (*β* = −0.043, 95% CI [−0.064, −0.021], *p* = 3.60 × 10*^−^*^4^), neutral (*β* = −0.031, 95% CI [−0.052, −0.010], *p* = 0.0045), and unpleasant (*β* = −0.023, 95% CI [−0.046, −8.82 × 10*^−^*^5^], *p* = 0.049). Together, these effects indicate that negative emotional intensity robustly modulates amygdala responses after accounting for baseline firing and between-subject variability. For hippocampal neurons selective for very pleasant ratings (Fig. 2f), the overall main effect of valence was also statistically significant (*F* (4, 3038.7) = 3.697, *p* = 0.0052), while baseline firing rate again strongly predicted task-evoked firing (*β* = −0.093, 95% CI [−0.130, −0.056], *p* = 6.70 × 10*^−^*^7^). The firing rates of these neurons in the very pleasant condition were higher than those for pleasant (*β* = −0.024, 95% CI [−0.037, −0.010], *p* = 0.002), and neutral stimuli (*β* = −0.023, 95% CI [−0.035, −0.009], *p* = 0.002), and marginally higher than those for very unpleasant stimuli (*β* = −0.016, 95% CI [−0.032, −8.10 × 10*^−^*^4^], *p* = 0.052) with no significant effect to unpleasant conditions (*β* = −0.011, 95% CI [−0.027, 0.0039], *p* = 0.14). These findings indicate that hippocampal neurons responsive to positive valence differentiate very pleasant stimuli from several, but not all, other emotional conditions.

To further examine valence-selective neuronal populations, we performed complementary analyses across pleasant-, neutral-, and unpleasant-selective neurons in both the amygdala and hippocampus (Fig. S3). The results show that pleasant- and neutral-selective neurons in both regions did not differentiate their preferred condition from other emotional conditions in both regions (*F* (4, 1723.3) = 2.082, *p* = 0.081 for pleasant, *F* (4, 1812.8) = 0.716, *p* = 0.581 for neutral in the amygdala; *F* (4, 2638.3) = 0.681, *p* = 0.605 for pleasant, *F* (4, 2167.3) = 0.339, *p* = 0.851 for neutral in the hippocampus). Neurons selective for unpleasant ratings exhibited significantly elevated firing specifically during unpleasant trials compared to all other valence categories in the hippocampus (*F* (4, 2042.2) = 4.562, *p* = 0.001; *β* = −0.024, 95% CI [−0.043, −0.006], *p* = 0.011 to very pleasant, *β* = −0.031, 95% CI [−0.049, −0.014], *p* = 9.6 × 10*^−^*^4^ to pleasant, *β* = −0.033, 95% CI [−0.051, −0.017], *p* = 2.5 × 10*^−^*^4^ to neutral, and *β* = −0.021, 95% CI [−0.042, −0.0004], *p* = 0.0541 to very unpleasant) but not in the amygdala (*F* (4, 2000.6) = 0.397, *p* = 0.232). These results suggest a regional dissociation in valence coding, with the amygdala preferentially responding to very unpleasant stimuli, and the hippocampus comprising multiple condition-specific neuronal subpopulations that selectively differentiate certain positive and negative emotional states.

### 2.3 From sustained power to transient beta bursts-uncovering emotion-specific dynamics

Having established the existence of valence-specific neuronal populations, we next examined whether neural oscillatory dynamics coordinated these responses. Time-frequency decomposition revealed stimulus-evoked changes—decreased low frequency power from theta band (4-8 Hz) to beta band (13-30 Hz) and increased gamma power (*>* 40 Hz) increased following image onset (cluster-based permutation, *p <* 0.05; Fig. 3a-b, top). However, these power modulations were identical across emotional conditions in both the amygdala and hippocampus (ART ANOVA, all *p >* 0.05; Fig. 3a-b, bottom), indicating that sustained beta and gamma power alone do not differentiate subjective emotional appraisal in our task.

**Fig. 3:**
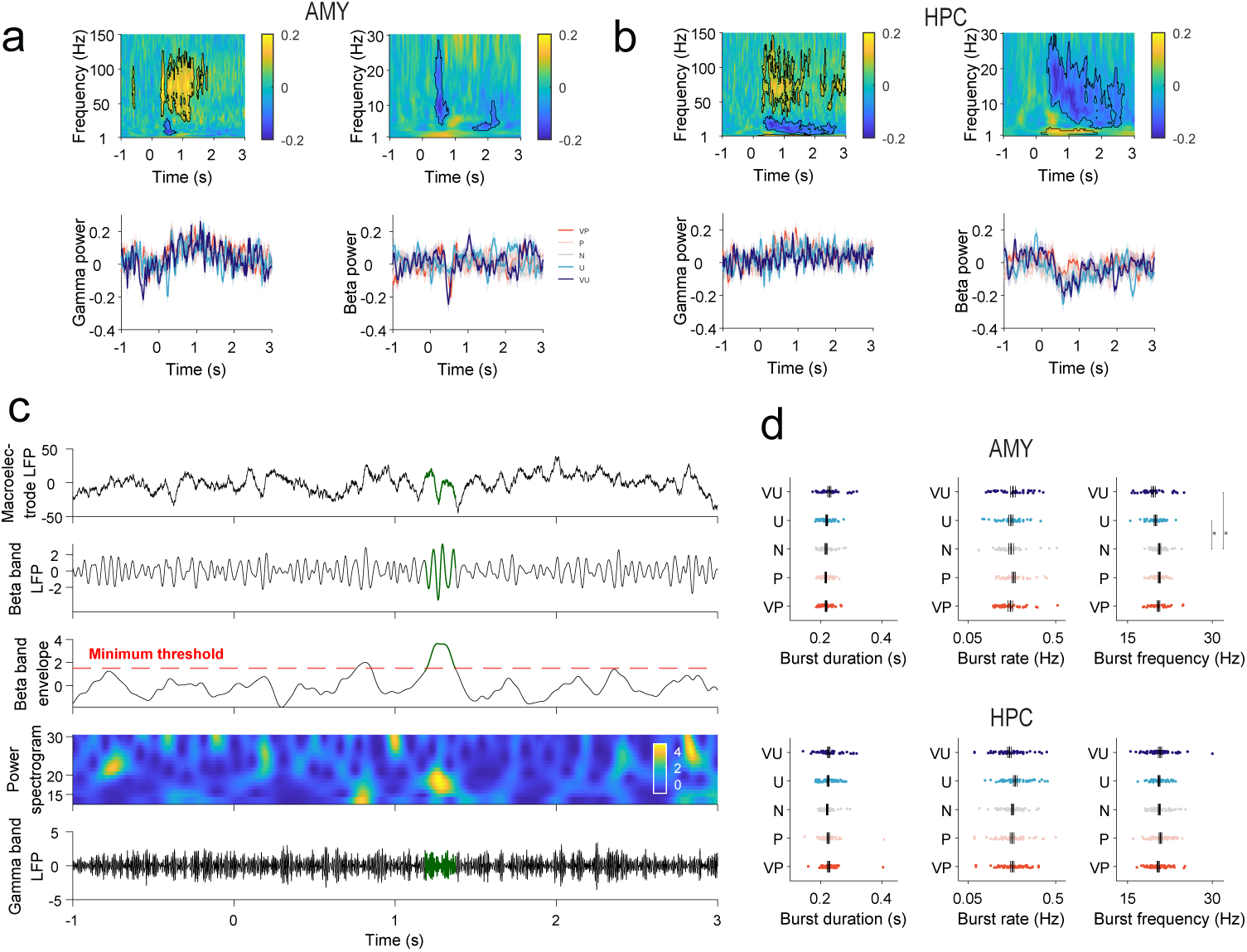
From sustained power to transient bursts-uncovering emotion-specific dynamics. **a**, Time-frequency analysis in AMY. Top: spectrograms showing stimulus-evoked changes in gamma (40-140 Hz, left) and beta (13-30 Hz, right) power. Black contours indicate significant changes from baseline (cluster-based permutation, *p <* 0.05). Bottom: averaged power traces for each valence rating (1-5) show no emotion-specific differences despite clear stimulus responses. (ART ANOVA FDR corrected *p >* 0.05). Shading: ± s.e.m. **b**, Same analysis for HPC, similarly showing no valence-specific power modulation. **c**, Beta burst detection pipeline. Example trial showing (top to bottom): raw macroelectrode LFP trace, LFP bandpass-filtered in the beta band, smoothed amplitude envelope of the beta-filtered signal with detection threshold (red dashed line), time-frequency spectrogram, and aligned gamma-band activity. Green highlighted line indicates detected bursts. **d**, Beta burst characteristics across emotional ratings. Top row: AMY; bottom row: HPC. Left to right: burst duration, rate, and peak frequency. Each dot represents one LFP channel; bars show group means. Asterisks indicate significant differences unpleasant vs neutral ratings in the AMY (\**p <* 0.05, permutation test with ART ANOVA multiple comparisons FDR corrected).

Additionally, to empirically validate our focus on the beta band—given the prominence of theta and alpha rhythms in prior emotional learning literature—we performed several control analyses. First, we conducted a 1/f-corrected spectral analysis using the IRASA (Irregular-Resampling Auto-Spectral Analysis) [57] method to separate genuine oscillatory peaks from the aperiodic background. This analysis revealed a significant, distinct oscillatory peak in the beta frequency range (13–30 Hz) in both the amygdala and hippocampus, confirming that the observed beta activity represents a functional rhythm rather than a spectral artifact or harmonic (Fig. S4a). Next, we evaluated the functional relevance of these bands for our specific task and found that traditional spectral power in the theta and alpha bands did not significantly differentiate between valence conditions. (Fig. S4b). The results establish beta activity as a genuine oscillatory component and indicate that sustained theta and alpha power do not account for valence-dependent effects.

We subsequently analyzed transient oscillatory beta burst dynamics [40, 58, 59]. First, we extracted brief high-power beta bursts (see example burst in Fig. 3c) from macroelectrode LFP recordings in both regions. Across all participants, we identified 13,260 bursts (5,097 in amygdala, 8,163 in hippocampus). In the amygdala, burst duration and rate were not significantly modulated by valence (all *p >* 0.05). However, peak frequency differed significantly across emotional ratings (*F* (4, 180) = 4.21*, p* = 0.0028). Stimuli rated as neutral (20.83 ± 1.36 Hz) were associated with higher peak frequencies than unpleasant ones (unpleasant: 19.83 ± 1.52 Hz, *p* = 0.037, very unpleasant: 19.65 ± 2.22 Hz, *p* = 0.009), with no other pairwise differences surviving after multiple comparisons correction using FDR. In the hippocampus, burst duration, rate, and peak frequency showed no significant main effect of valence (all *p >* 0.05). These results suggest that while the occurrence and duration of beta bursts were insensitive to valence, spectral characteristics in the amygdala may encode specific aspects of emotional appraisal.

To assess the dominance of bursting activity, we then compared burst duration across bands using a rhythm detection method adapted from Maher et al. [60]. and found that beta bursts occupied a significantly higher proportion of the signal duration than theta and alpha bursts (Fig. S4c). Together, these control analyses provide convergent evidence that valence-dependent dynamics may be selectively expressed in the beta band.

### 2.4 Amygdala beta bursts are associated with reduced hippocampal activity during negative emotions

To better understand the function of beta bursts, we examined their impact on local activity the amygdala and hippocampus separately, as well as on cross-regional neural dynamics, by analyzing high-gamma (HG) amplitude, which closely tracks local spiking activity, and single-neuron spiking activity during bursts. In both regions, beta bursts triggered robust high-gamma increases (80-140 Hz) compared to surrogate events (*p <* 0.05; Fig. 4a-b, top). We then asked whether HG amplitude was associated with valence ratings. In the amygdala, there was no significant change in burst-associated HG amplitude across emotional conditions (Fig. 4a, bottom). However, the very pleasant condition exhibited a higher HG compared to moderate conditions following bursts in the hippocampus (ART ANOVA followed by cluster based permutation testing, *p <* 0.05). This finding is consistent with increased firing rates of very pleasant-responsive neurons in the hippocampus. Together, these results suggest that beta bursts may play a role in coordinating the activity of local neuronal populations.

**Fig. 4:**
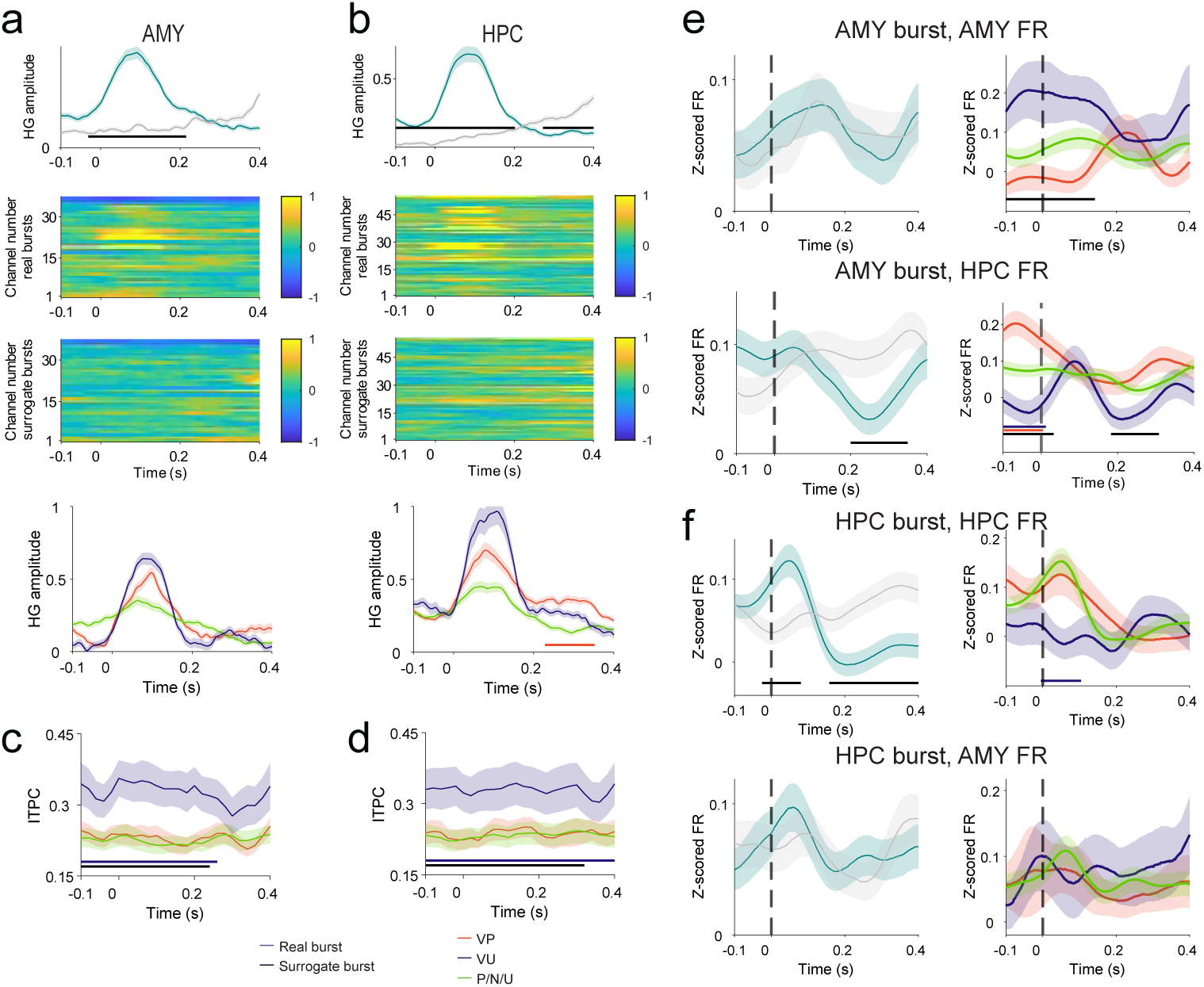
Asymmetric beta burst control: amygdala gates hippocampal emotional processing. **a**, AMY beta bursts are associated with local high-gamma (80-140 Hz) increase. Top: averaged HG amplitude for real bursts (purple) significantly exceeds surrogate events (black) (*p <* 0.05). Middle: single-channel heatmaps showing consistent HG enhancement across recording sites. Bottom: Comparisons of HG amplitude during bursts in the AMY between extreme and moderate emotional ratings. 0 is burst onset **b**, Same analysis for HPC beta bursts, showing similar local HG enhancement. **c,d**, Inter-trial phase coherence (ITPC) during beta bursts in AMY (c) and HPC (d). Very unpleasant stimuli (purple) show significantly stronger intertrial phase coherence than neutral and pleasant ratings (*p <* 0.05). **e**, Population neuronal response in the AMY and HPC during AMY bursts. Left panels: Comparisons of FRs in AMY neurons (n = 82 neurons) (top) and in HPC neurons (bottom, n = 136 neurons) aligned to AMY burst onset vs surrogate bursts. Only the HPC neurons show a decrease in FRs (cluster permutation test *p <* 0.05). Right panels: Increased AMY firing for very unpleasant ratings at the onset of AMY beta burst, while decreased HPC firing for very unpleasant ratings during AMY bursts (all, ART ANOVA FDR corrected *p <* 0.05, permutation test *p <* 0.05) **f**, Population neuronal response in the HPC and AMY during HPC bursts (n=136 neurons). Left panels: Comparisons of FRs in HPC neurons (top) and in AMY neurons (bottom) aligned to HPC bursts vs surrogate bursts. Only the HPC neurons show a decrease in FRs (cluster permutation test *p <* 0.05). Right panels: Decreased HPC firing for very unpleasant ratings at the onset of HPC beta burst (ART ANOVA FDR corrected *p <* 0.05, permutation test *p <* 0.05) but no effect on AMY firing pattern during HPC bursts (ART ANOVA FDR adjusted *p >* 0.05), indicating unidirectional AMY→HPC influence. Shading: ± s.e.m. Horizontal bars indicate significant periods (black: ratings VP vs VU; red: VP vs P/N/U; blue: VU vs P/N/U; all *p <* 0.05).

This increase in gamma during bursts coincided with spiking activity changes in the hippocampus, but not in the amygdala. Specifically, amygdala neurons did not show significant changes in firing rates after the onset of their own bursts (Fig. 4e, top left; *p >* 0.05), while hippocampal neurons did exhibit suppression of their firing rate after the onset of their own bursts. (Fig. 4f, top left; *p <* 0.05). However, this burst-firing relationship was emotion-specific. During very unpleasant ratings, amygdala neurons showed enhanced preburst activity followed by suppression during amygdala bursts (Fig. 4e, top right; *p <* 0.05). Hippocampal neurons exhibited earlier, deeper inhibition to very unpleasant ratings around hippocampus bursts(Fig. 4f, top right; *p <* 0.05). This pattern may reflect selective modulation of different neuronal subpopulations in each region.

Furthermore, we observed an asymmetric, valence-dependent pattern of cross-regional interaction during bursts: amygdala beta bursts were associated with a potent 80.96% suppression of hippocampal firing, with the maximal effect observed during very unpleasant ratings (Fig. 4e, bottom; *p <* 0.05). In contrast, hippocampal bursts exerted no reciprocal influence on amygdala single-neuron activity (Fig. 4f, bottom; *p >* 0.05).

To resolve whether the observed burst-related firing changes arose from specific neuronal classes, we decomposed firing-rate modulation during bursts by putative cell type in each region. This analysis builds on cell-type classification as shown in Fig. 5a,b and Fig. S6, which integrates waveform duration, firing rate, and autocorrelogram features, exhibits a bimodal trend in trough-to-peak durations, and yields consistent cell-type assignments across a broad range of classification thresholds. In the hippocampus, putative pyramidal neurons drove the firing rate decrease observed during very unpleasant ratings while narrow and wide putative hippocampal interneurons did not exhibit any statistically significant firing rate changes across valence conditions (Fig. 5c; *p <* 0.05). Further, amygdala beta bursts were followed by a suppression of hippocampal interneuron activity (narrow and wide) during very unpleasant trials, but not the activity of pyramidal cells (Fig. 5d; *p <* 0.05). We did not have enough amygdala interneurons (n=4) to conduct the converse analysis. These results suggest that this asymmetric relationship may reflect a hierarchical mechanism in emotional processing: the amygdala may modulate hippocampal processing specifically during very negative emotional states.

**Fig. 5:**
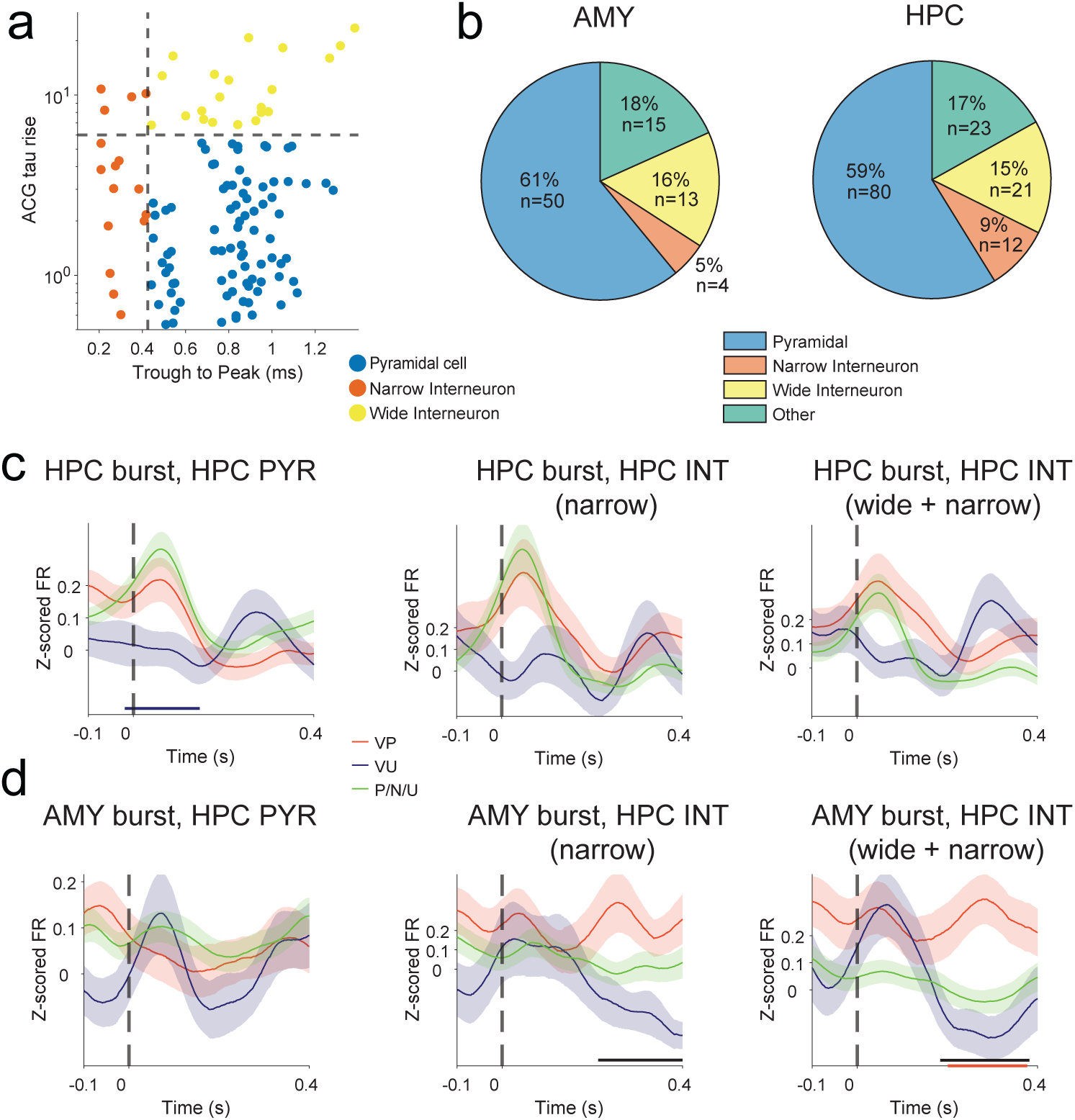
Hippocampal pyramidal cells (PYR) and interneurons (INT) show differing responses to beta bursts during negative emotional valence rating. **a**, Putative neuronal cell-type classification with autocorrelogram (ACG) tau rise time versus spike trough-to-peak duration for hippocampal single units, classified as putative pyramidal cells (blue), narrow interneurons (orange), and wide interneurons (yellow). Dashed lines indicate classification thresholds. **b**, Cell-type classification based on waveform characteristics. Pyramidal cells (AMY: 61% N=50, HPC: 59% N=80) constitute the majority, with narrow interneurons (AMY: 5% N=4, HPC: 9% N=12), wide interneurons (AMY: 16% N=13, HPC: 15% n=21), and unclassified cells making up the remainder. **c**, Cell-type-specific burst modulation in HPC neurons. Pyramidal cells (PYR, left, N=80) show early suppression during beta bursts for very unpleasant stimuli. In contrast, narrow interneurons (INT, middle, N=12) and wide interneurons (INT, right, N=21) did not exhibit significant difference across valence. **d**, Cell-type-specific AMY burst modulation of HPC neurons. Pyramidal cells (PYR, left, N=80) show no difference across valence. However narrow interneurons (INT, middle, N=12) and wide interneurons (INT, right, N=21) show suppression for very unpleasant stimuli (rating 5). This opposing pattern suggests beta bursts may coordinate emotional processing through differential modulation of excitatory and inhibitory circuits. Black bars: *p <* 0.05, cluster-based permutation test.

To characterize the directionality of amygdala–hippocampal interactions during beta bursts, we analyzed effective connectivity using the Phase Slope Index (PSI). The analysis revealed a robust, unidirectional hierarchy in which the amygdala consistently drives the hippocampus. Specifically, in the AMY→HPC direction during bursts in both regions, we observed significant positive PSI values(Fig. S5a, c, top) with peaks at the beta band (17.46 Hz), indicating that the amygdala leads. Conversely, in the HPC→AMY direction during bursts in both regions, significant negative PSI values were observed (Fig. S5b, d, top), further confirming that the amygdala leads the hippocampus. Importantly, PSI reflects the relative temporal lead–lag relationship between signals during beta-band activity and does not imply that beta bursts in one region trigger bursts in the other. Unlike firing rate modulation, which was valence-specific, the magnitude of this directional effect did not differ significantly across valence conditions (Fig. S5, bottom panels), suggesting that beta bursts support a stable amygdala-to-hippocampus communication channel regardless of valence.

### 2.5 Beta-gamma coupling distinguishes emotional valence through phase-specific modulation

Despite the absence of differences in spectral power between emotional ratings, we found that discrete, time-locked beta bursts modulate neuronal activity within and across regions during emotional processing. To further investigate whether phase-specific neural modulation could account for differences in emotional appraisal at the LFP level, we first analyzed ITPC during bursts. We found that the timing and magnitude of burst-locked dynamics varied with emotional valence: very unpleasant stimuli elicited stronger beta phase-locking in both the hippocampus and the amygdala, as indicated by increased ITPC (ART ANOVA test with multiple comparisons FDR correction *p <* 0.05; pairwise cluster-based permutation t test, *p <* 0.05; Fig. 4c-d).

Next, we quantified phase-amplitude coupling (PAC) between beta phase and gamma amplitude using a modulation index (MI) approach [61, 62]. While sustained beta power showed no differences across emotional conditions, PAC revealed robust emotion-specific modulation (Fig. 6a). The beta phase at which gamma amplitude peaked differed across emotional ratings (pairwise Mardia-Watson-Wheeler tests with multiple comparisons FDR correction, *p <* 0.01 for all conditions, Fig. 6b) with each emotional condition exhibiting a preferred beta phase (Rayleigh test, *p <* 0.01 for all conditions) in both regions. Notably, PAC strength, as measured by MI, was highest during very unpleasant trials in both the amygdala and hippocampus (Fig. 6c)and this emotion-specific coupling decreased systematically across the beta band (ART ANOVA test with FDR correction *p <* 0.05; pairwise cluster-based permutation t test, *p <* 0.05; Fig. 6d). These findings suggest that emotional valence may be encoded not only by the precise temporal coordination between oscillatory rhythms (PAC), but also by the strength of this coupling (MI).

**Fig. 6:**
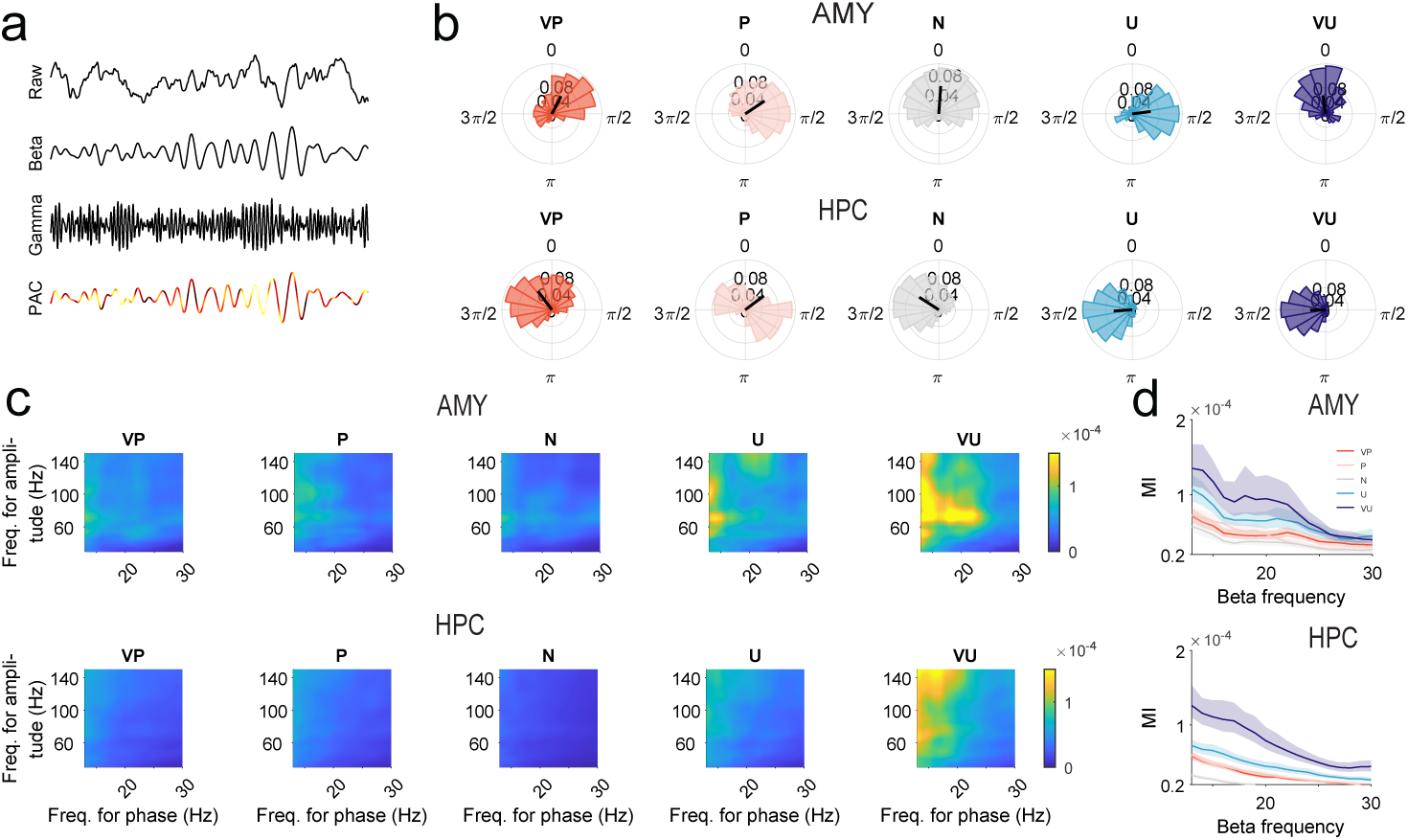
Beta-gamma PAC encodes emotional valence through coupling strength relationships. **a**, Example PAC analysis from a single trial. From top: raw LFP, beta-filtered (13-30 Hz), gamma-filtered (40-140 Hz), and computed PAC showing moments of strong coupling (red). **b**, Preferred coupling phase varies with emotional valence. Circular histograms showing the preferred beta phase at which gamma amplitude peaks for each emotional rating in the AMY (top) and HPC (bottom). All pairwise comparisons: *p <* 0.01, Mardia-Watson-Wheeler test with FDR correction. **c**, Emotion-specific PAC patterns. Comodulograms show coupling strength (MI) between beta phase and gamma amplitude frequencies for each emotional condition. Very unpleasant stimuli show strongest coupling at beta band in both AMY (top) and HPC (bottom). **d**, Group-averaged MI as a function of beta frequency for each emotional condition. Very unpleasant (purple) shows significantly stronger coupling than all other conditions. Shading area: ± s.e.m.

### 2.6 Effect of high arousal on single-unit firing and LFP response

To evaluate the effect of arousal on valence-related neural responses, we re-analyzed key LFP and single-unit data by restricting the Very Pleasant condition to trials with the top 30% standardized arousal ratings. This was performed across all subjects using the IAPS + NAPS dataset. We also performed the same analysis in a subset of subjects with IAPS-only stimuli, where arousal levels for Very Pleasant trials more closely matched those of Very Unpleasant trials.

The single-unit results remained consistent (Fig. S7a-b). Specifically, amygdala beta bursts were associated with decreased firing of hippocampal neurons during Very Unpleasant trials compared to the highest-arousal Very Pleasant trials. However, the highest-arousal Very Pleasant trials exhibited significantly increased inter-trial phase coherence (ITPC), higher than that of the Very Unpleasant and moderate trials (Fig. S7c-f). This increased ITPC and was not observed when Very Pleasant trials were collapsed regardless of arousal (Fig. 4).

To better understand the influence of arousal on ITPC, we next compared the top 30% arousal ratings of very pleasant and very unpleasant trials (Fig. S7g-h). Here we observed significantly increased ITPC in both extreme valence conditions compared to the moderate ratings but there was no longer any significant ITPC differences between Very Pleasant and Very Unpleasant trials. This suggests that that ITPC is influenced and varies with high arousal arousal.

## 3 Discussion

Our results suggest a new neural mechanism for human emotional processing: transient beta bursts that temporally correlate with amygdala-hippocampus communication through millisecond-precision temporal dynamics. We identified a subset of neurons in both the amygdala and hippocampus that displayed responsiveness to emotional stimuli, with firing rates differing across subjective emotional valence ratings. While the average spectral power in the beta and gamma bands did not discriminate among emotional conditions, beta burst dynamics in the amygdala and hippocampus revealed robust emotion-specific modulation across scales—from single-neuron firing to cross-regional communication. Furthermore, emotional valence may be encoded through systematic phase relationships rather than through power changes, suggesting a temporal code for emotion. Importantly, these findings provide evidence of coordinated neural dynamics rather than a definitive mechanistic account of inhibitory gating. Determining the circuit mechanisms underlying these dynamics will require causal perturbations and computational modeling.

We believe that our findings contribute to affective neuroscience by providing rarely available concurrent recordings of single-unit and oscillatory activities in the amygdala and hippocampus. Previous works suggest that the amygdala rapidly encodes emotional valence [63–65]. Consistent with this, we identified amygdala neurons sensitive to very unpleasant stimuli, approximately 500-800 ms post-stimuli, replicating prior intracranial studies with a similar time window (e.g., [66, 67]). Hippocampal neurons, which were mostly from the anterior hippocampus, also showed valence sensitivity to both positive and negative stimuli, aligning with existing evidence that the ventral hippocampus contributes to emotion in rodents [18, 68]. This dual selectivity in the hippocampus may reflect a contextual role of encoding, whereas the strongest aversive signals were preferentially expressed in the amygdala. The latency of hippocampal neuronal responses was similar to that of the amygdala, and is consistent with other literature examining human amygdala and hippocampus during emotional perception tasks. For instance, Wang et al., 2014 [66] showed that amygadala neurons responsive to faces with negative or positive emotions exhibited significant differences in firing rate around 625-ms post-stimulus-onset. In Mormann et al., 2008 [69], average response latency in the hippocampus was around 394 ms, and those in the amygdala 397 ms, at which time around 50% of the responsive units exhibited significant differences in firing rate from baseline. Response latencies reported in other human regions span a broad range, including approximately 600–820 ms in ventromedial prefrontal and anterior cingulate cortex [70] and 300–520 ms in regions such as dorsal striatum [71], placing the 500–800 ms latencies observed here in amygdala and hippocampus well within reported range.

Beyond spiking activity, oscillatory dynamics also play a role: prior studies have linked beta [17, 72] and gamma [51, 73] oscillations to emotional evaluation and salience encoding. Our time-frequency decomposition revealed similar spectral features, however, these were insufficient to distinguish emotional valence appraisals, echoing concerns that sustained oscillations may overlook key dynamic features of emotional processing [74, 75]. Transient beta bursts offered a more precise neural marker. This aligns with recent paradigm in a rodent study [18], suggesting that transient beta bursts may reflect a key neurocomputational mechanism for emotion processing.

Our findings may also offer new insight into the functional role of beta bursts in affective encoding within the human medial temporal lobe. Emerging evidence suggests that oscillatory bursts—rather than sustained rhythms—play a key role in modulating cognitive and emotional function [49, 76]. Beta bursts, in particular, have been implicated in routing and gating information during cognitive tasks [40, 77]. Rodent and nonhuman primate studies show that beta bursts within hippocampus and amygdala circuits regulate attention states [41] and memory encoding [18]. Extending this, we found that beta bursts in the amygdala and hippocampus were modulated by emotional stimuli and linked to transient changes in spiking activity and gamma amplitude. These bursts occurred concurrently during affective information processing, especially for highly pleasant or unpleasant stimuli. Neuronal responsiveness was stronger during burst windows, suggesting that beta bursts may coordinate ensemble-level encoding of emotional valence. Moreover, amygdala bursts exhibited a different peak frequency for neutral versus negative valence ratings, while the hippocampus lacked such valence-specific burst modulation (Fig. 3d). This suggests that there might be differences in regional specialization across the hippocampus and amygdala. Beta bursts also showed robust cross-frequency coupling (CFC) with gamma, supporting hierarchical organization across frequency bands during emotion-related tasks [18, 40, 42, 49].

The amygdala and hippocampus form a tightly interconnected circuit critical for emotional processing and affective memory. Anatomically, these regions share reciprocal projections: the basolateral amygdala projects to dorsal and ventral hippocampus [17, 63], while the hippocampus sends input back via the subiculum [43]. Functionally, this circuitry is crucial for integrating emotional information with contextual memory, enabling behavioral responses and long-term memory encoding. For example, fMRI studies show increased amygdala-hippocampus activation during emotional memory and aversive recall [78–80]. Meanwhile, human electrophysiology confirms dynamic coordination between these regions during emotion processing. Zheng et al. [75] demonstrated theta–alpha multiplexing between amygdala and hippocampus, revealing frequency-specific communication for affective salience. Costa et al. [73] also reported enhanced synchrony of the cross-regional theta phase with gamma activity during aversive memory tasks. While these studies established theta and alpha synchrony as channels for information transfer supporting memory encoding, our results suggest that beta bursts are associated with inhibitory dynamics during rapid appraisal, with beta burst coordination temporally linked to amygdala-hippocampus communication and interneuron activity [18]. We observed that beta bursts not only modulated local gamma amplitude and neuronal firing within the amygdala and hippocampus but also exerted cross-structural effects. Specifically, beta bursts of the amygdala appear to modulate the firing rates in the ipsilateral hippocampus, but not vice versa. Cell-type analysis further supported this circuit understanding: that the suppressed firing rate seen in the hippocampus during the amygdala burst was driven by hippocampal interneurons and not by pyramidal cells. (Fig. 5d), This finding aligns with rodent studies showing that somatostatin interneurons modulate synchronization between these two regions [18]. The asymmetric, burst-triggered coordination supports a dynamic, state-dependent model of emotional coding, where in transient oscillatory events shape communication between amygdala and hippocampus [17, 81].

Our ITPC and PAC analyses suggest that phase relationships may be better than oscillatory power in distinguishing emotional states. Both the amygdala and hippocampus exhibited enhanced inter trial phase coherence (Fig. 4c-d) specifically for very unpleasant valence ratings. Furthermore, while the preferred phase for maximal gamma amplitude varied across valence conditions, the coupling strength was remarkably specific with very negative/high arousal valence ratings showing increased coupling strength. Together, these results suggest that phase modulation is associated with selective enhancement and temporal coordination of affective information processing, especially for strong negative emotions.

Aberrant amygdala-hippocampus dynamics have been implicated in psychiatric conditions such as major depressive disorder, PTSD, and generalized anxiety [64, 65, 82, 83]. Disruptions in oscillatory coordination—particularly in beta and gamma bands—and cross-regional communication may contribute to maladaptive emotional encoding. Our findings suggest a potential mechanism for emotional processing that may inform future research on neuropsychiatric interventions. Specifically, beta bursts in amygdala and hippocampus, by modulating local spiking and gamma activity, may serve as real-time physiological markers of emotional states. This temporally precise neural motif is especially relevant for closed-loop neuromodulation strategies, including deep brain stimulation, transcranial alternating current stimulation, and transcranial magnetic stimulation [84].

Rodent studies have shown that manipulating somatostatin interneurons during beta coherence bursts can influence anxiety-related behavior [18]. In humans, high-frequency activity (HFA)—a proxy for local spiking [85] in MTL—has been used to modulate emotional recall [20] and alleviate depressive symptoms via personalized closed-loop stimulation [86]. By identifying amygdala beta bursts as a potentially modifiable behaviorally relevant neural event, our results suggest that this transient, dynamic view of emotional processing may open new avenues for understanding, and perhaps modulating, disorders of emotional regulation

Several limitations merit consideration. First, recordings from epilepsy patients may not generalize to healthy populations. Our participant cohort has a heterogeneous clinical background, encompassing different psychiatric and mood disorder histories as well as treatment with anti-seizure medications. Second, while our task effectively elicited emotional responses, naturalistic emotional experiences involve additional complexity that may not be captured here. Third, the correlational nature of our findings precludes causal inference—whether beta bursts actively shape emotional processing remains unresolved without direct manipulations such as burst-inducing stimulation as mentioned above. Fourth, while normative self-report ratings confirmed that the very unpleasant stimuli were associated with higher arousal, this was not seen in eye-tracking measures. This discrepancy likely stems from the short stimulus duration (3 s) in our task, as prior literature indicates that while pupil responses to emotional stimuli can exhibit early transient changes [87, 88], arousal-mediated differentiation is most consistently observed in later autonomic windows (e.g., 2–6 s in [89] or 2-7 s in [90]), beyond the early response phase. Notably, some studies report (e.g., [91])) that pupil dilation is greatest within early windows (1–4 s) but does not vary by valence during this period. Together, these findings indicate that our data do not rule out arousal-related confounds in our neural findings. Additionally, we focused on beta bursts due to their alignment with emotional stimuli, transient activity in other bands and their interaction with beta, such as through cross-frequency coupling may also contribute to emotional processing. Although our results implicate beta bursts as a key dynamic coordinating amygdala–hippocampal activity, establishing beta bursts as a distinct mechanistic unit will ultimately require generative circuit models and causal manipulations, which remain important directions for future work. Lastly, we acknowledge that our study is purely mechanistic, and thus any claims as to its therapeutic implication remain tentative. Future studies should employ causal interventions, include psychiatric clinical populations, incorporate more comprehensive autonomic measures, and examine multi-band burst interactions. Nevertheless, by revealing how transient beta bursts correlate with emotional processing through asymmetric dynamics and phase-specific encoding, our findings establish a new framework for understanding, and potentially treating, emotional processing and dysfunction.

## 4 Methods

### 4.1 Human participants

Fifteen participants (9 female, 6 male; age: 34.13 ± 12.15 years) with pharmacologically intractable epilepsy were enrolled at Yale New Haven Hospital. All procedures were approved by the Yale University Institutional Review Board and participants provided with written informed consent. Recordings occurred during clinical monitoring with depth electrodes placed solely for clinical purposes. Channels with epileptiform activity (IED) were excluded using automated IED detection and manual inspection. Data within 4 hours of clinical or electrographic seizures were excluded. Clinical characteristics (Table S1), including age, handedness, sex, anti-seizure medications, Beck Anxiety Inventory (BAI, [92])and Beck Depression Inventory (BDI, [93]) scores, are provided in Supplementary Table 1. BAI and BDI scores were obtained from each participant’s routine presurgical neuropsychological assessment. Detailed demographic and clinical information is provided in Table S2. All patients had medically intractable epilepsy. Specific epilepsy etiologies and current psychiatric medications are listed and no patients in this cohort had radiological evidence of mesial temporal sclerosis (MTS, Table S2).

### 4.2 Electrophysiological recording

Participants underwent stereotactic implantation of Behnke-Fried depth electrodes (Ad-Tech Medical) containing eight 40-µm microwires protruding 3-4 mm from the macroelectrode tip, enabling simultaneous recording of LFPs and single-unit activity from the amygdala and hippocampus (Fig. 1a). Microwire signals were digitized at 30 kHz (0.3-7.5 kHz bandpass) using a NeuroPort system (Blackrock Microsystems). Macroelectrode signals were sampled at 2 kHz via the clinical EEG system [94, 95].

### 4.3 Electrode localization & selection

Pre-implantation T1-weighted MRI scans were co-registered with post-implantation CT scans. Electrode positions were identified using LeGUI [96] and mapped to standardized Montreal Neurological Institute (MNI) space (Table S3). Gray and white matter boundaries were automatically segmented to exclude contacts residing in white matter. Regional assignments used the Neuromorphometrics atlas [97]. For visualization, we plotted the MNI coordinates of all macroelectrodes across participants on the 152 template brain from the MNI [98, 99].

For inter-regional analyses, we included only the subset of patients who had simultaneous intracranial electrode coverage in both the amygdala and hippocampus. Analyses were performed strictly within hemispheres (ipsilateral connections). To avoid selection bias in subjects with multiple contacts per region, we did not select a single representative channel; instead, we chose all possible ipsilateral Amygdala-Hippocampus channel pairs and averaged the results for the subject.

### 4.4 Behavioral task

Participants viewed 120 color photographs while seated in hospital beds. Visual stimuli were presented on a 24-inch monitor (2K, 120 Hz) using MATLAB and Psychtoolbox. Latencies between stimulus onsets and TTL pulses were logged and later compensated for in neural analyses. Participants were provided with a Gamepad or keyboard for ratings. Each trial consisted of fixation (1 s), image presentation (3 s), and rating period (maximum 5 s). The participants rated the images from 1 (very pleasant) to 5 (very unpleasant), with moderate values corresponding to pleasant (2), neutral (3), and unpleasant (4) (Fig. 1b). Trials with incorrect or absent responses were excluded from behavioral and neural analyzes. The stimuli consisted of emotionally evocative images selected from two validated datasets: the International Affective Picture System (IAPS) for 8 patients and the Nencki Affective Picture System (NAPS) for 7 patients, which was used to match visual properties across categories (e.g., luminance, contrast, spatial frequency). Both datasets included an equal number of images from four semantic categories: scenes, objects, animals, and humans with 30 images per category, ensuring consistent visual diversity and balanced emotional content. All images were presented at their original size and were novel to the participants at the time of testing. Verbal instructions were given prior to the task. Eye tracking was performed in 12 sessions from 6 healthy subjects and 3 epilepsy patients using Tobii Pro (60 Hz) to examine arousal differences across valence conditions.

### 4.5 Behavioral analysis

We quantified the distribution of emotional ratings across all trials and subjects (Fig. 1c). The trials assigned to each category were counted and differences between rating conditions were tested using ART ANOVA with permutation tests and FDR correction (Fig. S1a). Response times (RT) was defined as the interval between image offset and the participant’s rating response by to button press (Fig. 1d and Fig. S1b). Trials with RTs shorter than 0.2 s (anticipatory responses) or longer than 5 s (inattentive responses) were excluded, resulting in the removal of 8.87% of trials. ART ANOVA test was also performed followed by post-hoc permutation with FDR correction across ratings and image categories. For pupil analysis (Fig. 1e and Fig. S1c), raw pupil diameter data were preprocessed to extract the most valid eye per sample. Based on the standard preprocessing pipeline for Pupil Core data, samples with low confidence (*<* 0.8) were discarded. Artifacts (*>* 2.5*SD* change) were removed and missing samples interpolated. Data were smoothed with a third-order Butterworth low-pass filter (4 Hz low-pass), and z-score normalized per subject. ART ANOVA assessed differences across emotional ratings, with pairwise cluster-based permutation t-test for time-resolved analysis. To validate the arousal content of our stimuli independent of physiological measures, we also analyzed the standardized mean arousal scores from the IAPS and NAPS normative datasets for all images presented(Fig. S1d-e). We compared these normative scores across our five valence categories using ART ANOVA. We further analyzed pupil responses using a basis-function general linear model adapted from previous work (e.g., Yebra et al., 2019 [100]). Each trial-wise pupil time course was modeled as a linear combination of an early light-reflex component and a later cognitive component, each represented by an Erlang gamma function with the same estimated parameters. To directly test for potential arousal-related confounds, trials were grouped based on standardized arousal ratings (high vs low arousal), and the cognitive component was compared between these conditions at the subject level. To quantify evidence for the absence of arousal effects, Bayes Factors [101] were computed for the paired comparison between high- and low-arousal conditions.

### 4.6 LFP preprocessing

Macroelectrode channels were selected based on their anatomical localization in the hippocampus and amygdala. LFPs were downsampled to 1000 Hz and common-average referenced across all channels within each subject. Bipolar referencing between neighboring contact pairs was used for beta burst detection [102, 103]. All prepreprocessing was conducted with the FieldTrip toolbox[104]. Data were bandpass filtered with a zero-phase lag FIR filter (1-200 Hz) and notch filtered at 60 Hz harmonics (59–61 Hz, 119–121 Hz, 179–181 Hz). Epochs were extracted from -1 to 3s relative to stimulus onset. Artifacts were rejected using automated detection (z *>* 3), followed by manual inspection to remove segments containing non-physiological artifacts (e.g., movement, clipping, or sudden transients).

### 4.7 Power and Time-frequency analysis

To verify the presence of distinct oscillatory peaks above the aperiodic background, we employed the Irregular-Resampling Auto-Spectral Analysis (IRASA) method [57]. This technique separates the power spectral density (PSD) into fractal (1/f) and oscillatory components by resampling the time-series data at non-integer factors and taking the geometric mean of the resulting PSDs. We analyzed the baseline-corrected LFP data to quantify the significant power deviations in the beta band (13–30 Hz) relative to the underlying fractal component.

Time-frequency decomposition was performed to characterize task-induced power changes relative to baseline activity for each channel based on FieldTrip toolbox with Morlet wavelets (4-14 cycles) over 1 to 150 Hz in 1 Hz steps [105]. Power was baseline-normalized using the -1 to 0 s pre-stimulus window. Cluster-based permutation tests were used to assess differences in spectral power between the pre-stimulus and stimulus windows (0–3 s after image onset) or across valence conditions. Statistical significance was determined by identifying clusters in time-frequency space exceeding chance levels based on the permutation distribution.

### 4.8 Beta burst detection and characterization

Beta bursts were identified and characterized based on established protocols from recent intracranial and MEG literature [40, 58, 59]. Signals were bandpass filtered (13–30 Hz; two-pass, fourth-order Butterworth) and transformed via the Hilbert method to extract amplitude envelopes, smoothed with a 20 ms moving average (Fig. 3c). Beta bursts detection criteria includes: (1) power spectrum *>* 2 SD above the trial mean, (2) ≥ 3 oscillatory cycles (i.e., three peaks and three troughs), (3) smoothened envelope *>* 1.5 SD above the tiral mean, and (4) duration between 80 ms to 500 ms. Note that the first two criteria were applied on the bandpass-filtered signal, while the last two criteria used the envelope. Manual verification by two independent raters was used for burst verification based on comparative assessments (10% of detected bursts, achieving above 90% agreement) of signal quality. Sensitivity analysis confirmed results were robust to detection threshold variations (1.5-2.5 SD). The characteristics (Fig. 3d), including burst rate, duration, and peak frequency, were calculated per trial. Peak frequency was calculated using a wavelet-based spectral estimate (cfg.toi = -1 all:3). Burst rate was defined as the number of bursts per 3 s stimulus period normalized by trial length. ART ANOVA with permutation t tests were used for analysis between conditions in these two regions.

Meanwhile, we quantified the prevalence of bursts across theta (4–8 Hz), alpha (8–12 Hz), and beta (13–30 Hz) bands using the eBOSC (extended Better Oscillation Detection) framework, following the protocol of Maher et al. [60]. For each frequency band, we calculated the “burst duration proportion,” defined as the percentage of the total task duration that the LFP signal exceeded the amplitude threshold (95th percentile) for a minimum of 3 cycles. This metric provided a standardized measure of burst density to compare the dominance of transient oscillatory events across frequency bands.

### 4.9 PAC

The instantaneous amplitude of HG and phase of beta of the LFPs for each electrode during the 3 s following image onset of each trial were extracted using the Hilbert transform (Fig. 6). Phases were binned into 18 bins of 20 degrees. PAC was quantified by the MI, using the Kullback-Leibler distance method combined with Shannon’s entropy [61, 62]. Significance was assessed against 100 surrogate MI distributions generated by trial shuffling. All comparisons of MI used ART ANOVA test followed by permutation tests with FDR correction (*p <* 0.05). Pairwise Mardia–Watson–Wheeler tests with FDR correction were used to compare the phase preference of the maximum amplitudes across emotional ratings, with the Rayleigh test used to compare the phase distribution against a uniform distribution.

### 4.10 Spike sorting

Microwire signals were bandpass filtered (300-3000 Hz) and spike-sorted using Wave Clus3 (single-cell quality metrics are shown in Fig. S2) [106]. Detected spike clusters were subsequently curated manually using custom MATLAB scripts and the Wave Clus GUI. Units were retained if: (1) *<*5% inter-spike intervals (ISIs) violations (ISI *<* 3 ms), (2) *>*200 spikes per session, and (3) physiological waveforms [58, 107]. Waveform stability and isolation quality were visually verified through ISI histograms, PCA projections, and consistency across time. This yielded 136 hippocampal and 82 amygdala units across participants. Neurons were classified as putative pyramidal cells or narrow interneurons based on waveform characteristics [108]. Wide interneurons were identified by trough-to-peak duration *>*0.425 ms, acg tau rise *>* 6*ms*, and firing rates *<*10 Hz, while narrow interneurons had short trough-to-peak duration (*<*0.425 ms) and higher firing rates (*>*10 Hz). The remaining cells with wide waveforms and low firing rates are assigned as pyramidal cells. The metrics were selected based on extensive literature on electrophysiology recordings of different cell types in medial temporal lobe [109–111]. Notably, the waveform and autocorrelogram firing properties of wide interneurons do not clearly separate from the clusters of pyramidal or narrow interneurons so we placed them as a separate category. Spike trough-to-peak durations exhibited a bimodal pattern in the histogram. Hartigan’s Dip Test yielded a dip statistic of 0.0439 with a p-value of 0.0579. The trend toward bimodality may reflect noise and variability of human intracranial recordings.

### 4.11 Neural response analysis

Firing rates were calculated in 2-ms non-overlapping bins and smoothened with a Gaussian kernel (σ = 200 ms). Population peristimulus time histograms (PSTHs) were computed by pooling neurons within each region for each emotion rating condition. Cluster-based permutation tests identified emotion-specific responses. Responsive neurons (Fig. 2a, b) were defined as units that showed significant rate changes during 300-1300 ms after image onset versus baseline (-1000-0 ms) (two-tailed cluster-based permutation t-test, *p <* 0.05) [112, 113]. This yielded multiple subsets of neurons preferentially responding to specific emotional rating levels. To assess population-level dynamics (Fig. 2c, d), emotional-responsive units from all subjects were pooled separately for the hippocampus and amygdala to form regional populations. For each brain region and each emotional rating, we computed trial-averaged population firing rates. These population PSTHs were then grouped and compared across different emotional conditions using contrasts that were independent of the unit-selection step. Group-level statistical comparisons were conducted using a cluster-based permutation test (*p <* 0.05).

To evaluate how emotional valence modulated neuronal firing in the amygdala and hippocampus while accounting for subject-level variability, we fitted a population-level linear mixed-effects model (LME) [52, 114] to all trials from valence-selective neurons in Fig. 2e-f. For each trial, the dependent variable was the mean firing rate during the task window (FR_task_; 0.5–1.0 s after stimulus onset). Baseline firing rate (FR_baseline_; –1.0 to 0 s) was included as a covariate to adjust for baseline-dependent variability across units, and Valence was modeled as a categorical fixed effect (5 levels). To account for the hierarchical structure of the data (multiple neurons recorded within each patient), subject and unit were modeled as random intercepts:

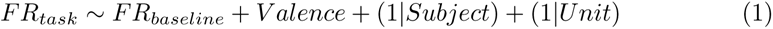

The reference category for Valence was set to the lowest valence level (Valence = 1). Fixed-effect coefficients reflect adjusted differences in task-period firing relative to this reference at the mean baseline firing rate. Degrees of freedom were estimated using the Satterthwaite approximation. Post-hoc contrasts and 95% confidence intervals were computed from fixed-effects estimates. The same procedure was applied to very pleasant -responsive neurons in hippocampus, with very pleasant as the reference. *β* coefficients indicate adjusted differences from the reference valence, and model-predicted FRs were computed at the mean baseline FR. Significance for contrasts with the reference was taken from LME coefficient *p*-values with FDR correction. Model assumptions were assessed through visual inspection of residuals and random-effects distributions for the LME models.

### 4.12 LFP and spike activity during beta bursts

To examine the physiological relevance of transient beta bursts, we analyzed both the dynamics of HG amplitude (Fig. 4a, b) and neuronal firing rates (Fig. 4e, f) aligned to beta burst onset within these two regions. HG amplitude was obtained using a zero-phase FIR filter and Morlet wavele transform with a frequency resolution of 0.5 Hz (80:0.5:140 Hz) and a temporal resolution of 20 ms from -100 ms to 400 ms around burst start time. This window was selected based on the physiological properties of the detected beta events and to maximize statistical power. Across subjects and regions, the mean beta-burst duration was 218.2 ± 25.1 ms in the AMY and 223.0 ± 29.6 ms in the HPC (Fig. 3), corresponding to at least 3 oscillatory cycles at the observed beta frequencies. The chosen window therefore captures the pre-burst baseline, the full burst epoch, and the immediate post-burst recovery period [40, 44, 58]. The extracted gamma envelopes were z-score normalized within each trial relative to the baseline window (-1000-0 ms) and averaged per channel. Responses were then concatenated across subjects to form group-level representations. For neuronal spiking data, Gaussian-smoothed firing rates (200 ms kernel) were aligned to the onset each beta burst and aggregated across trials. Inter-regional analysis was conducted in subjects who have electrodes in both the HPC and AMY (N = 10) (Table S3). For each selected subject, we computed the inter-regional metric (e.g., burst-firing rate modulation) for each of the HPC–AMY channel pairs; these pairwise values were then z-scored at the subject level before group statistics to avoid bias from differing channel counts.

To determine the directionality of information flow during beta bursts, we calculated the Phase Slope Index (PSI) [43]. PSI estimates the causal direction of interaction by quantifying the consistency of the phase lag across a frequency band; a positive slope indicates that the signal from the first region leads the second. We calculated the “Relative Change in PSI” (Real - Surrogate) for four configurations: (a) AMY→HPC during AMY bursts, (b) HPC→AMY during AMY bursts, (c) AMY→HPC during HPC bursts, and (d) HPC→AMY during HPC bursts. Directionality was assessed by comparing the time-resolved PSI against zero (indicating significant directional flow) and comparing average PSI values across valence conditions.

To assess the specificity of bursts, we created surrogate bursts by selecting random time points within 200 ms before or after each real burst, excluding any intervals with artifacts or overlapping bursts. Surrogate-aligned HG amplitudes and firing rates were calculated using the same procedure. Subsequently, HG amplitude and Gaussian-smoothed firing rates were aligned to burst onset and compared to surrogate events [40]. We also computed inter-trial phase coherence (ITPC, Fig. 4c, d) of beta-band LFPs to assess whether beta bursts were associated with phase-aligned activity. Phase estimates were extracted using Morlet wavelets, and ITPC was calculated per channel and concatenated across subjects. Finally, to evaluate cross-regional interactions, we extracted high-gamma amplitude and firing rates from ipsilateral electrodes during beta bursts detected in the other region, aligned to burst onset times. The significance of real bursts versus surrogate bursts was assessed using cluster-based permutation tests. The comparisons across conditions during bursts were evaluated by ART ANOVA with multiple comparisons FDR correction and pairwise permutation tests.

### 4.13 Statistical analysis

We employed non-parametric statistical methods throughout to account for violations of normality and the presence of unequal trial counts across conditions. Statistical tests were chosen based on the number of groups being compared and whether the analysis was time-resolved. Non-parametric approaches were adopted for all analyses as described below:

To compare conditions without a temporal dimension, we used two-tailed permutation tests to assess differences in trial-level distributions between the two groups. To assess differences in time-resolved signals between two conditions, we used two-tailed cluster-based non-parametric permutation tests. For the permutation, we generated 1000 random surrogate datasets by shuffling the labels of two groups with the Monte Carlo method. At each timepoint, an uncorrected p-value was calculated; timepoints with *p <* 0.05 were grouped into contiguous clusters. The empirical p-values for each cluster were computed as the proportion of permutations yielding a cluster statistic greater than or equal to the observed one [105, 115].

For comparisons involving three or more conditions without a temporal dimension, we used an ART ANOVA test [116–121] as a non-parametric alternative to repeated-measures ANOVA. This method was selected to accommodate non-normal data distributions and unequal sample sizes across valence conditions while permitting the assessment of interaction effects. If a significant effect was detected, post-hoc pairwise comparisons were performed using permutation tests with p-values corrected for multiple comparisons using the Benjamini–Hochberg false discovery rate (FDR) procedure. For time-resolved multi-group analyses, ART ANOVA was performed at each time point, followed by pairwise comparison using cluster-based permutation test with FDR correction. All statistics were conducted in Matlab, except ART ANOVA which was finished in R.

## 5 Data availability

All data produced in this work are available on request from the corresponding author via email, with no conditions or restrictions.

## 6 Code availability

Custom MATLAB code associated with this study for data analysis are available online at https://github.com/damisahlab/AMY-HPC-Emotion.

## Acknowledgements

We thank the members of the Yale Comprehensive Epilepsy Center for their excellent patient care, the patients who participated in this study. This study was supported by grants from the National Institutes of Health (R01MH138291 ECD); The Hypothesis Fund Seed Grant (ECD), Center for Brain and Mind Health (ECD, AK, IHR, AS), VA National Center for PTSD and Wellcome Trust Grant: 310142/Z/24/Z (ECD, AK, IHR and JK).

## 7 Author Contributions

Conceptualization, E.C.D.; Methodology, E.C.D, Y.Z., Y.H., G.P., B.Z., A.A. and A.P.T; Software, Y.Z., Y.H., B.Z., A.A. and A.P.T; Validation, Y.Z., Y.H.; Formal analysis, E.C.D, Y.Z., Y.H.; Investigation, E.C.D, Y.Z., Y.H.; Resources, E.C.D.; Data curation, E.C.D, Y.Z., Y.H.; Writing—original draft, Y.Z., Y.H.; Writing—review and editing, E.C.D, Y.Z., Y.H., G.P., B.Z., A.A, A.P.T, A.S., Z.Z., S.O, I.H., A.K., J.K., K.S., X.G., C.P.; Visualization, Y.Z., Y.H.; Supervision, E.C.D.; Project administration, E.C.D, Y.Z., Y.H.; Funding acquisition E.C.D.

---

**Fig. S1:**
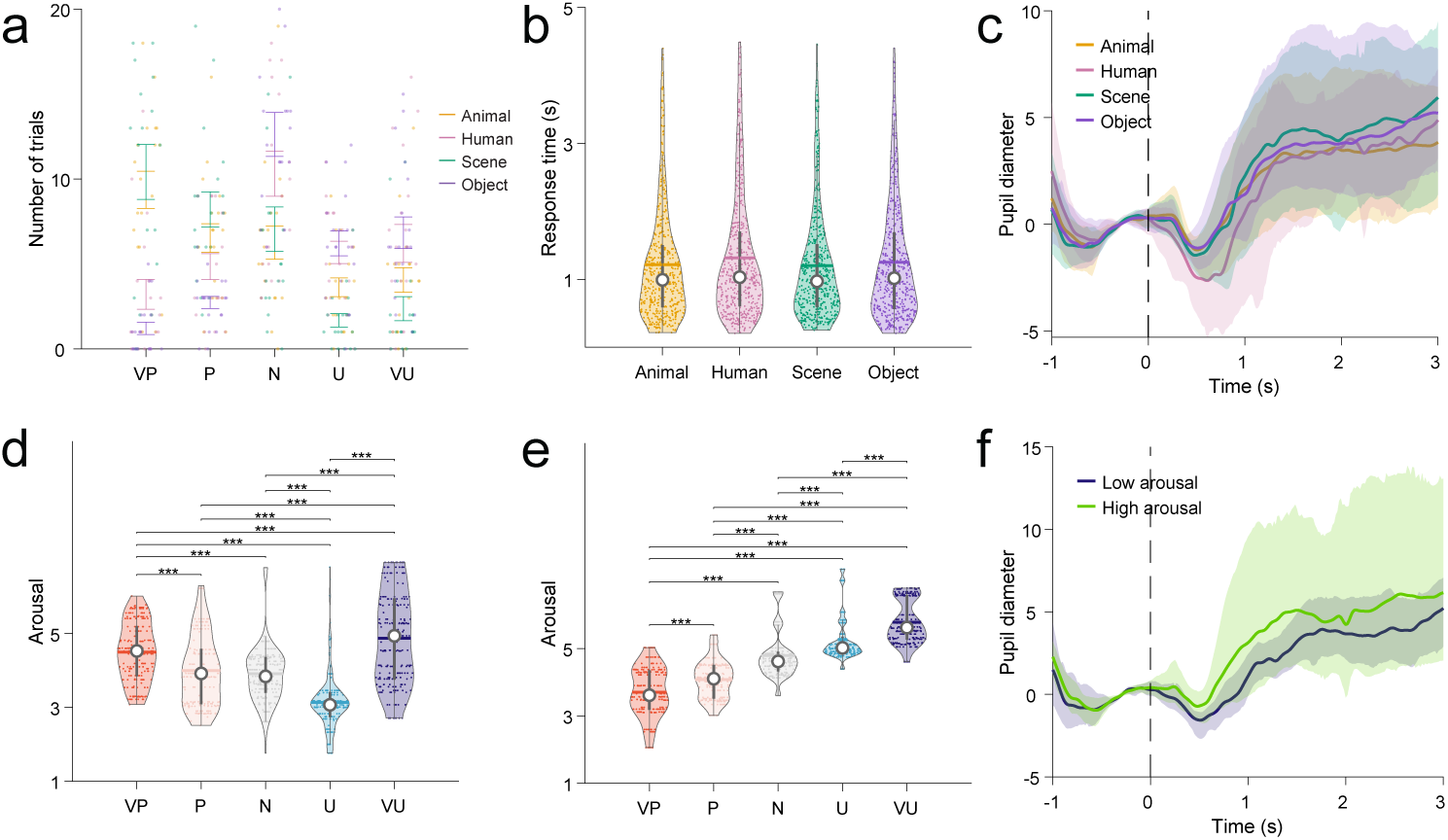
Stimulus category effects on behavior are limited to rating distributions. **a**, Distribution of emotional ratings varies by stimulus category. Objects and humans elicit predominantly neutral ratings, while animals and scenes evoke more extreme ratings, particularly pleasant. Bars show mean number of trials ± s.e.m. ART ANOVA followed by permutation tests with FDR correction revealed significant category effects (all *p <* 0.001). **b**, Response times show no category specificity. Mean RTs (± s.e.m.) are similar across all four stimulus categories (ART ANOVA: *F*_(3,2072)_ = 0.73, *p* = 0.53). **c**, Pupil diameter responses are category-independent. Time courses show no arousal differences between stimulus categories (ART ANOVA: *p >* 0.05). Shaded areas: ± s.e.m. Vertical dashed line indicates image onset. **d-e**, Distribution of standardized arousal scores derived from IAPS (**d**) and NAPS (**e**) normative databases) for the images selected for this study. ART ANOVA test followed by permutation tests with FDR correction. \*\*\**p <* 0.001. **f**, Pupil diameter responses across standardized ratings for low arousal and high arousal conditions.

**Fig. S2:**
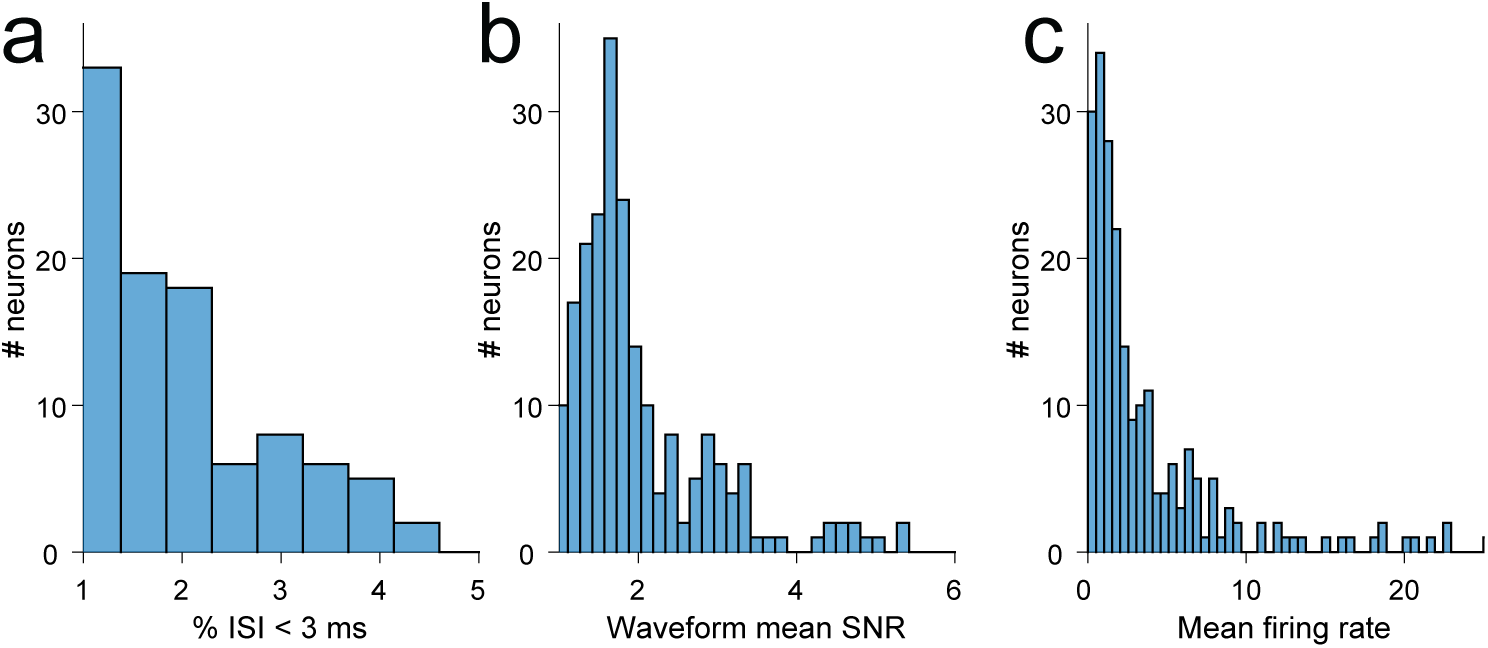
Quality metrics to evaluate reliable single-unit isolation. **(a)** Proportion of inter-spike intervals (ISI) below 3 ms (1.13% ± 1.63%, mean ± s.d.). **(b)** Signal-to-noise ratio (SNR) for the mean waveform across all spikes as compared to the standard deviation of the background noise (2.01 ± 1.00, mean ± s.d.). **(c)** Average firing rate within the entire recording session for all identified single units (3.77 ± 4.83, mean ± s.d.).

**Table S1:**
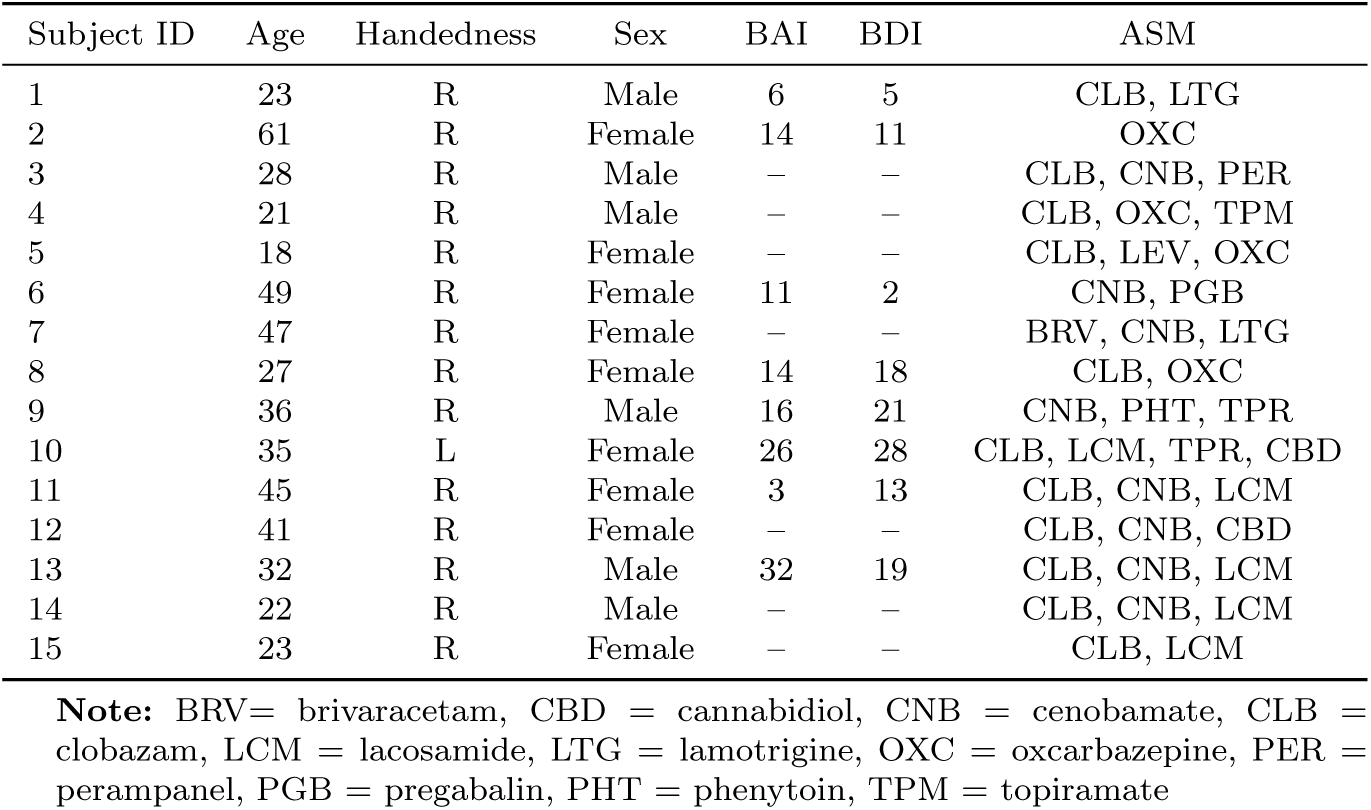
Subject information.

**Fig. S3:**
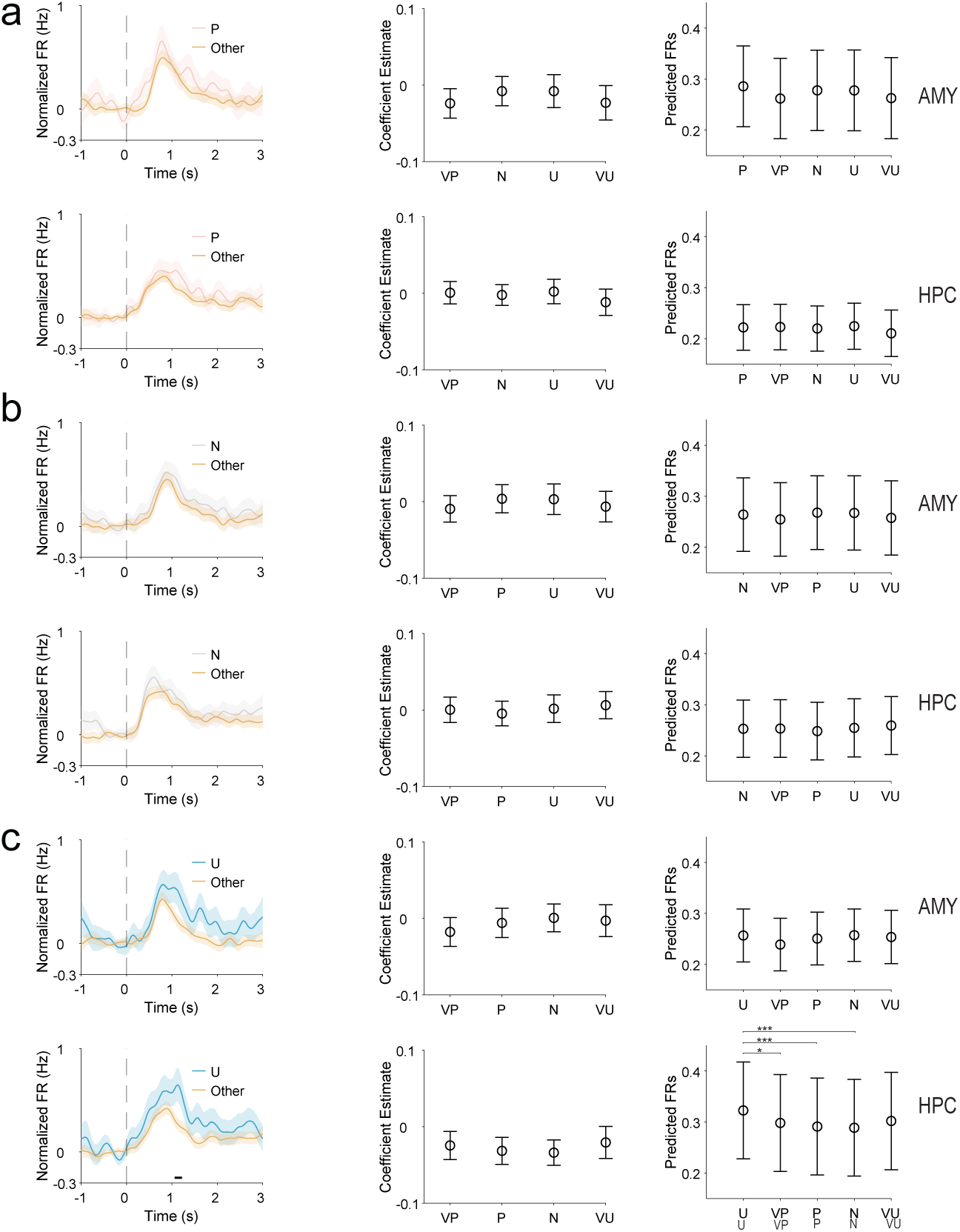
Valence-selective neuronal populations in amygdala and hippocampus. Population responses of neurons selective for pleasant (**a**), neutral (**b**), and unpleasant (**c**) emotional ratings in the amygdala (AMY; top row of each panel) and hippocampus (HPC; bottom row). Left panels show normalized firing rates aligned to image onset for trials corresponding to the preferred valence (colored) compared to all other trials (gray) within each selective population. Middle panels display coefficient estimates (±95% CI) from population-level LME controlling for baseline firing rate. Right panels show model-predicted firing rates (±95% CI) for each valence condition. \**p <* 0.05, \*\*\**p <* 0.001.

**Fig. S4:**
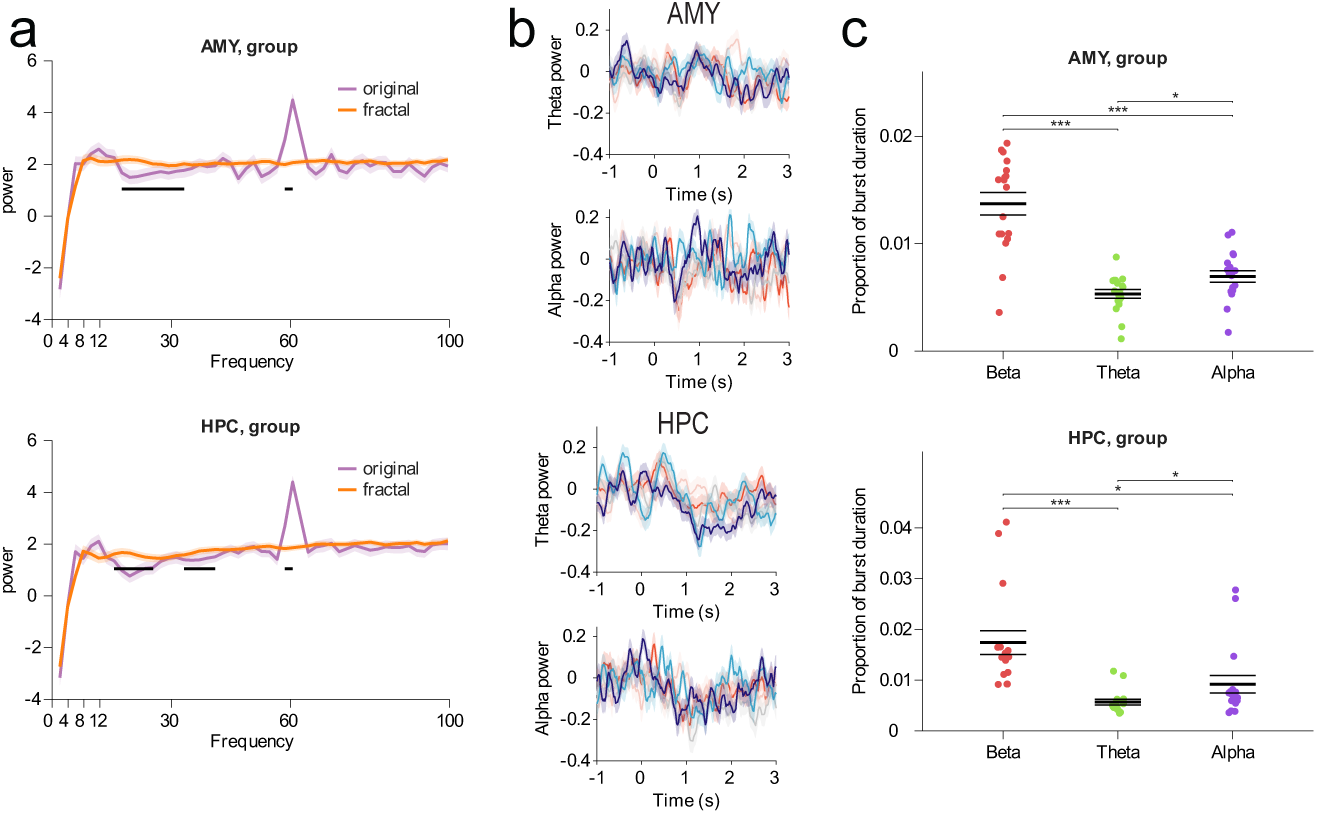
Validation of beta band dominance and specificity. **a**, 1/f-corrected spectral analysis using IRASA. The plot displays the oscillatory power spectrum (purple) separated from the fractal (1/f) background component (brown). A distinct, significant peak is observed in the beta band for both amygdala and hippocampus, confirming beta as a genuine oscillatory rhythm rather than a spectral artifact. **b**, Time-resolved power analysis for theta (left) and alpha (right) bands. Average power in these low-frequency bands did not significantly differentiate between emotional valence conditions in the AMY or HPC (VP vs. VU vs. N; *p >* 0.05) **c**, Comparison of burst prevalence across frequency bands. Bar plots show the mean proportion of total trial time occupied by bursts in the theta, alpha, and beta bands. Beta bursts exhibited a significantly higher proportion of burst duration compared to theta and alpha bursts (*p <* 0.05), indicating that beta bursting is the dominant transient oscillatory band during this task. Error bars indicate ± s.e.m.

**Fig. S5:**
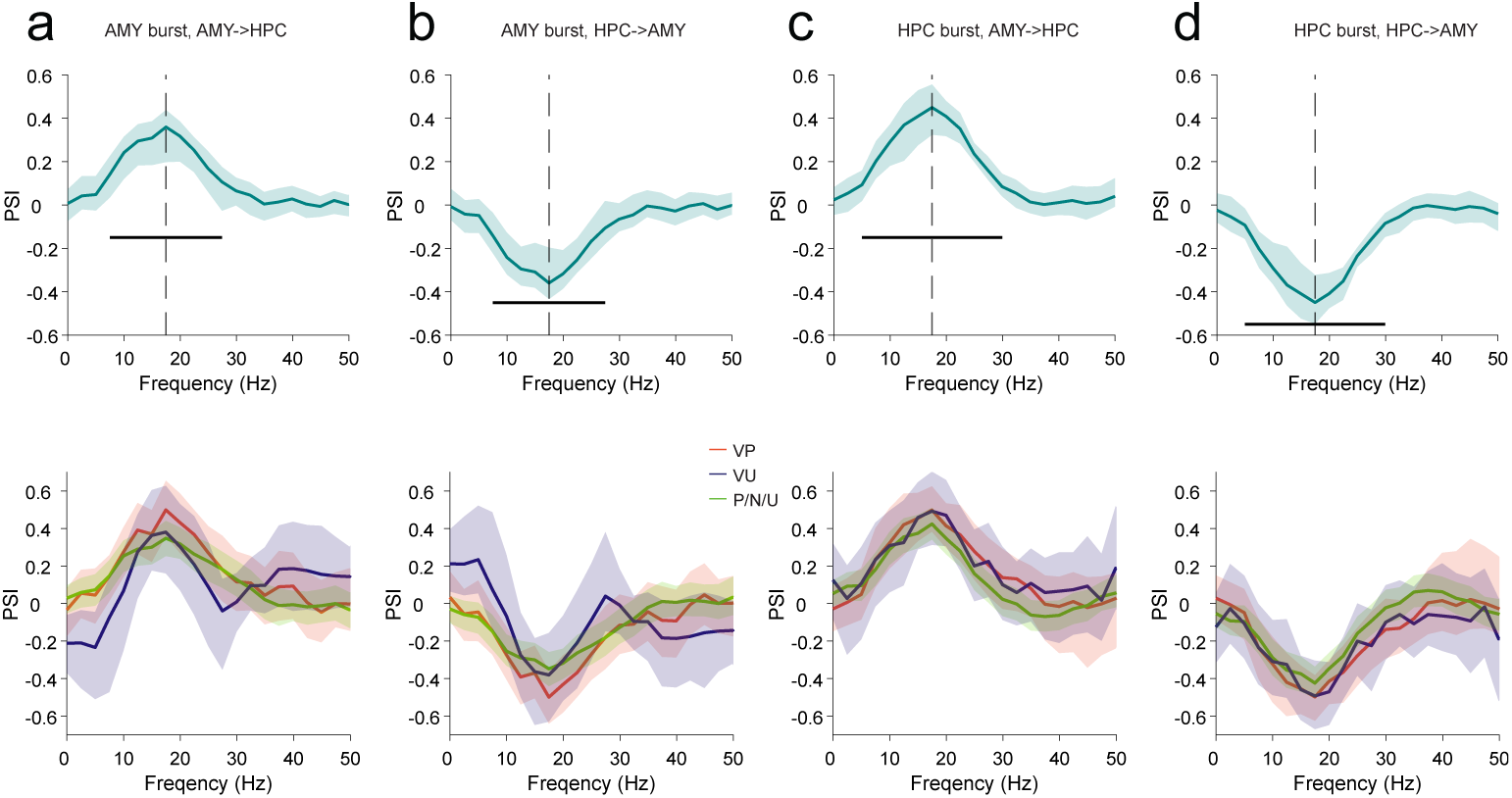
Unidirectional amygdala (AMY) → **hippocampus (HPC) drive during beta bursts.** The figure displays the relative change in PSI. Green lines indicate mean PSI; black bars indicate significance (*p <* 0.05). **a**, AMY→HPC PSI during AMY Bursts. The positive value suggests AMY leads HPC activity. **b**, HPC→AMY PSI during AMY Bursts. The negative value suggests AMY leads HPC activity (HPC lags). **c**, AMY→HPC PSI during HPC Bursts. The positive value suggests AMY leads neuronal activity, even during HPC bursts. **d**, HPC→AMY PSI during HPC Bursts. The negative value suggests AMY leads and HPC lags. Bottom rows in each panel show that the magnitude of this modulation is consistent across valence conditions (*p >* 0.05). Error bars: ± s.e.m.

**Fig. S6:**
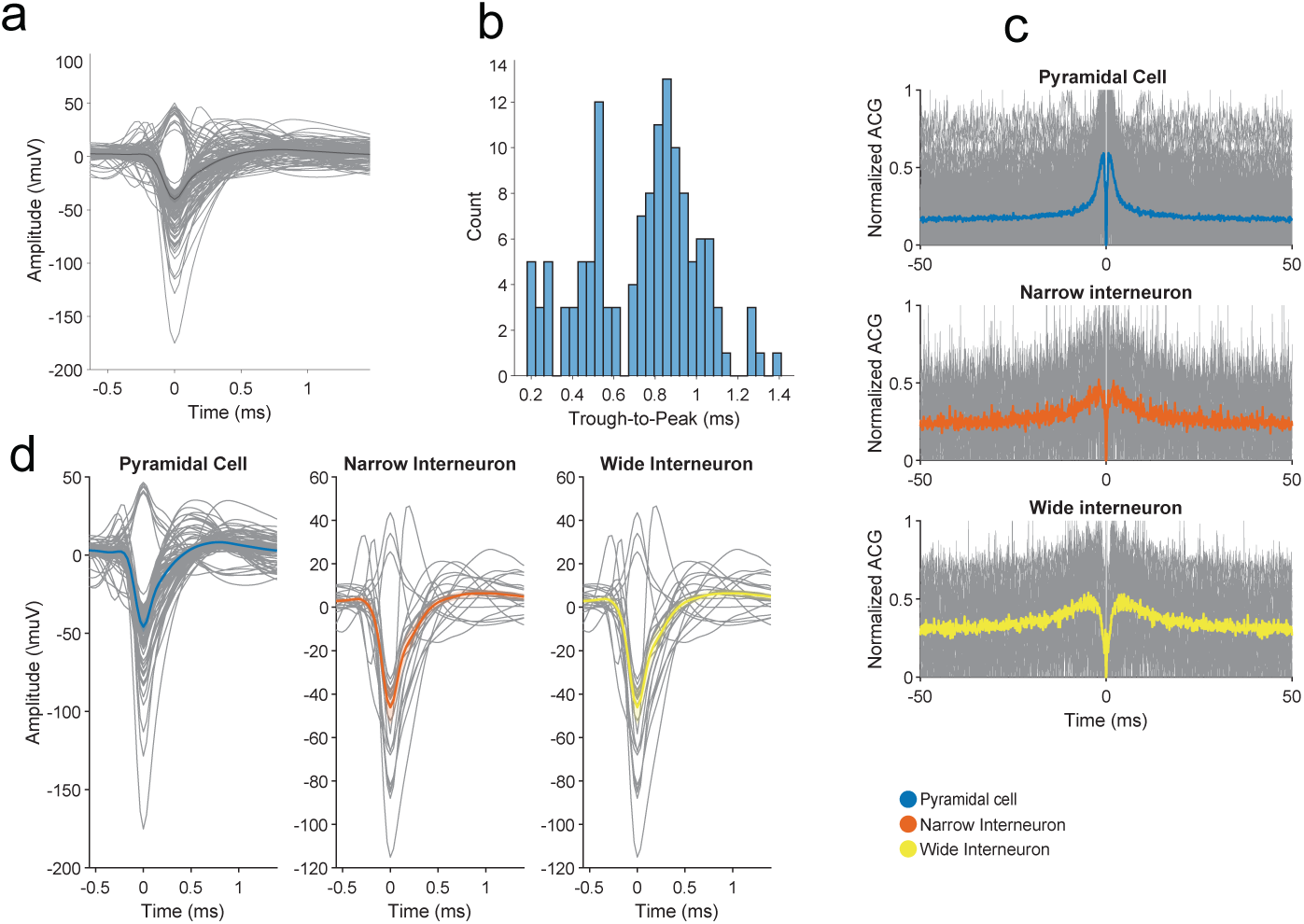
Putative neuronal cell-type validation. **a**, Mean-normalized spike waveforms from all units aligned to the trough. **b**, Histogram of spike trough-to-peak durations for putative pyramidal cells (blue), narrow interneurons (orange), and wide interneurons (yellow), showing a trend toward bimodality (Hartigan’s Dip Test: statistic = 0.0439, p = 0.0579). **c**, Average normalized autocorrelograms for each putative cell type (gray: individual units; color: mean). **d**, Average spike waveforms for putative pyramidal cells, narrow interneurons, and wide interneurons.

**Fig. S7:**
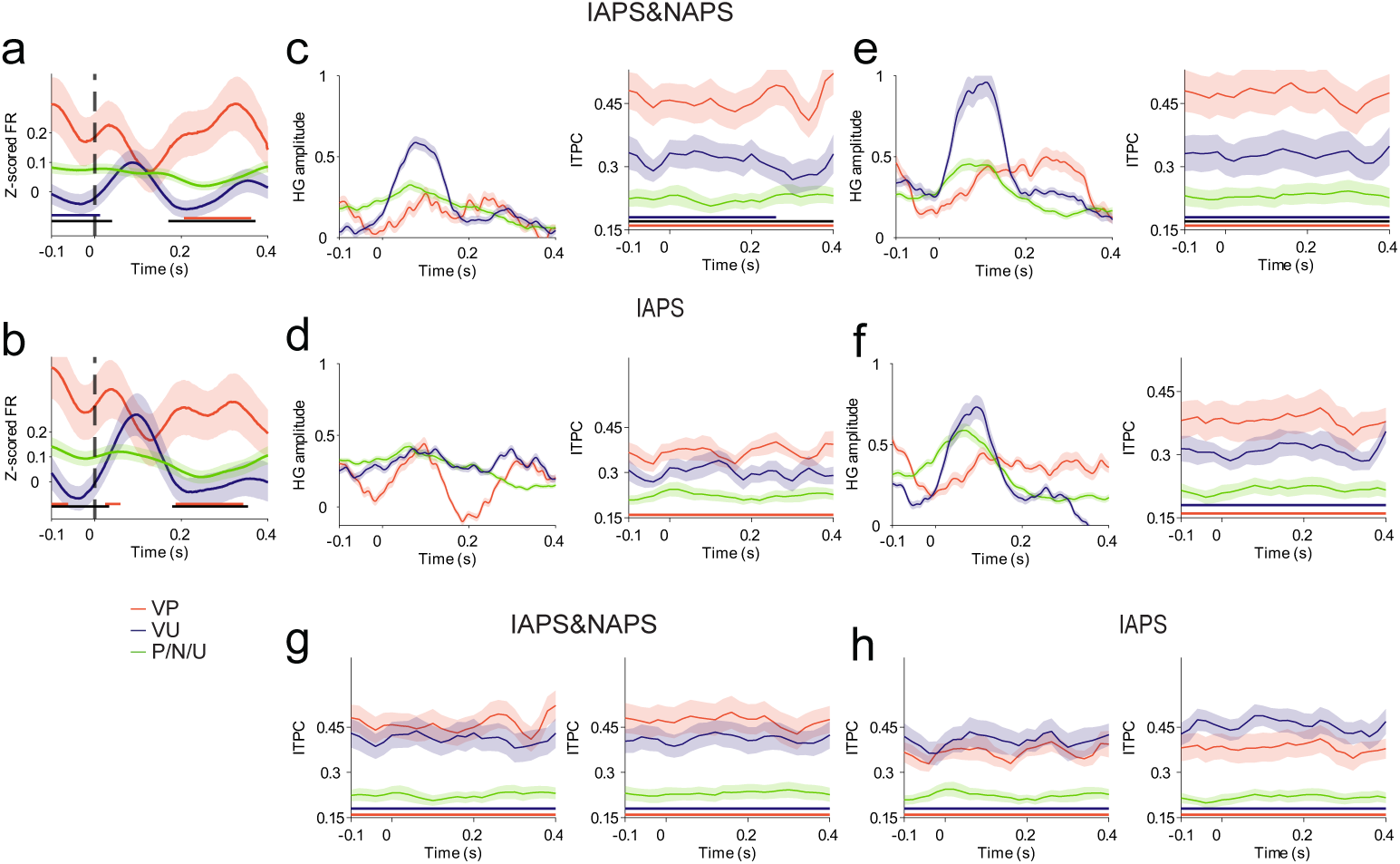
Effect of high arousal on single-unit firing and LFP response. Key single unit/LFP results after selecting top 30% arousal ratings for the very pleasant condition from all subjects (IAPS and NAPS datasets) (**a**, **c**, **e**) and with IAPS-only dataset in a subset of subjects (**b**, **d**, **f**). **a**, **b**, Population firing rates (z-scored) of HPC neurons aligned to AMY beta burst onset. **c**, **d**, HG amplitude (left) and ITPC (right) during beta bursts in the AMY. **e**, **f**, HG amplitude (left) and ITPC (right) during beta bursts in the HPC. **g**-**h**: ITPC (left: AMY; right: HPC) after selecting the top 30% arousal ratings for both very pleasant and very unpleasant trials from all subjects (IAPS and NAPS) (**g**) and IAPS only in a subset of subjects (**h**). Shaded areas indicate ± s.e.m. Horizontal bars indicate significant periods (black: ratings VP vs VU; red: VP vs P/N/U; blue: VU vs P/N/U; all *p <* 0.05).

**Table S2:** Patient Demographics and Clinical Characteristics.

| Subject ID | Epilepsy Etiology | Psychiatric Medications<sup>a</sup> | Epileptic Tissue<sup>b</sup> | Temporal Sclerosis |
| --- | --- | --- | --- | --- |
| 1 | Left frontal lobe cortical dysplasia | None | No | No |
| 2 | Bitemporal medial temporal lobe epilepsy; | None | Yes | No |
| 3 | Right frontal lobe polymicrogyria | Paroxetine | No | No |
| 4 | Right occipital lobe epilepsy | None | No | No |
| 5 | Right temporal neocortical lesion [temporal pole] | None | No | No |
| 6 | Right frontal lobe polymicrogyria [sylvian fissure] | None | No | No |
| 7 | Bitemporal anti-GAD encephalitis | None | Yes | No |
| 8 | Bitemporal epilepsy | None | Yes | No |
| 9 | Bilateral frontal lobe epilepsy | None | No | No |
| 10 | Right temporal lobe epilepsy | None | Yes | No |
| 11 | Bitemporal epilepsy; GAD65+ autoimmune | None | Yes | No |
| 12 | Bitemporal epilepsy; GAD65+ autoimmune | None | Yes | No |
| 13 | Temporal neocortex epilepsy; cortical dysplasia | Sertraline | No | No |
| 14 | Left temporal lobe epilepsy | Sertraline | Yes | No |
| 15 | Fronto-Temporal neocortex broad context | None | No | No |

Abbreviations: M, Male; F, Female; R, Right; L, Left; B, Bilateral; SOZ, Seizure Onset Zone.
<sup>a</sup> Psychiatric medications defined as agents prescribed specifically for mental health indications.
<sup>b</sup> Indicates if the analyzed Amygdala/Hippocampus electrodes were within the clinician-defined epileptic tissue.
**Note:** No patients in this cohort had Mesial Temporal Sclerosis (MTS).

**Table S3:** MNI coordinates of microwires for different brain regions.

| Electrode ID | Patient ID | Location | Side | X | Y | Z |
| --- | --- | --- | --- | --- | --- | --- |
| 1 | 1 | Hippocampus | Left | -25.68 | -18.76 | -19.12 |
| 2 | 1 | Amygdala | Left | -20.63 | -5.92 | -20.86 |
| 3 | 2 | Hippocampus | Left | -24.66 | -18.37 | -17.27 |
| 4 | 2 | Hippocampus | Right | 27.60 | -16.62 | -17.10 |
| 5 | 2 | Amygdala | Left | -18.97 | -6.17 | -20.79 |
| 6 | 2 | Amygdala | Right | 24.96 | -6.31 | -25.30 |
| 7 | 3 | Hippocampus | Right | 37.76 | -13.55 | -18.30 |
| 8 | 3 | Amygdala | Right | 16.08 | -2.12 | -22.97 |
| 9 | 4 | Hippocampus | Right | 20.56 | -15.02 | -15.85 |
| 10 | 5 | Hippocampus | Right | 27.65 | -15.71 | -23.18 |
| 11 | 5 | Amygdala | Right | 21.79 | -4.43 | -21.48 |
| 12 | 6 | Hippocampus | Right | 18.68 | -7.66 | -20.70 |
| 13 | 6 | Amygdala | Left | -18.22 | -9.02 | -25.21 |
| 14 | 7 | Amygdala | Right | 21.58 | -3.52 | -26.20 |
| 15 | 7 | Amygdala | Left | -20.98 | -7.06 | -23.74 |
| 16 | 8 | Hippocampus | Left | -19.09 | -7.05 | -25.35 |
| 17 | 8 | Hippocampus | Right | 23.33 | -12.54 | -20.00 |
| 18 | 8 | Amygdala | Left | -19.86 | 1.47 | -27.02 |
| 19 | 8 | Amygdala | Right | 19.99 | 0.05 | -28.69 |
| 20 | 9 | Hippocampus | Left | -26.11 | -9.49 | -23.71 |
| 21 | 9 | Hippocampus | Right | 24.24 | -10.36 | -19.10 |
| 22 | 10 | Hippocampus | Right | 25.41 | -12.53 | -17.62 |
| 23 | 10 | Hippocampus | Right | 31.39 | -28.20 | -6.98 |
| 24 | 10 | Hippocampus | Left | -28.83 | -21.17 | -13.09 |
| 25 | 10 | Amygdala | Right | 20.59 | -1.82 | -24.19 |
| 26 | 10 | Amygdala | Left | -20.49 | -5.15 | -21.68 |
| 27 | 11 | Hippocampus | Left | -23.11 | -14.84 | -15.39 |
| 28 | 11 | Hippocampus | Left | -22.35 | -31.47 | -4.94 |
| 29 | 11 | Hippocampus | Right | 26.51 | -15.82 | -15.76 |
| 30 | 11 | Amygdala | Left | -16.84 | -5.02 | -18.27 |
| 31 | 12 | Amygdala | Left | -15.93 | -5.97 | -24.35 |
| 32 | 12 | Hippocampus | Left | -21.00 | -16.91 | -19.73 |
| 33 | 12 | Hippocampus | Right | 21.70 | -23.40 | -21.56 |
| 34 | 13 | Hippocampus | Right | 25.22 | -9.19 | -20.95 |
| 35 | 13 | Amygdala | Left | 19.27 | -3.14 | -18.30 |
| 36 | 14 | Hippocampus | Left | -33.66 | -33.05 | -8.26 |
| 37 | 14 | Hippocampus | Left | -31.56 | -14.61 | -16.81 |
| 38 | 15 | Amygdala | Right | 19.80 | -1.10 | -20.07 |
| 39 | 15 | Hippocampus | Right | 33.51 | -14.97 | -17.82 |

**Table S4:**
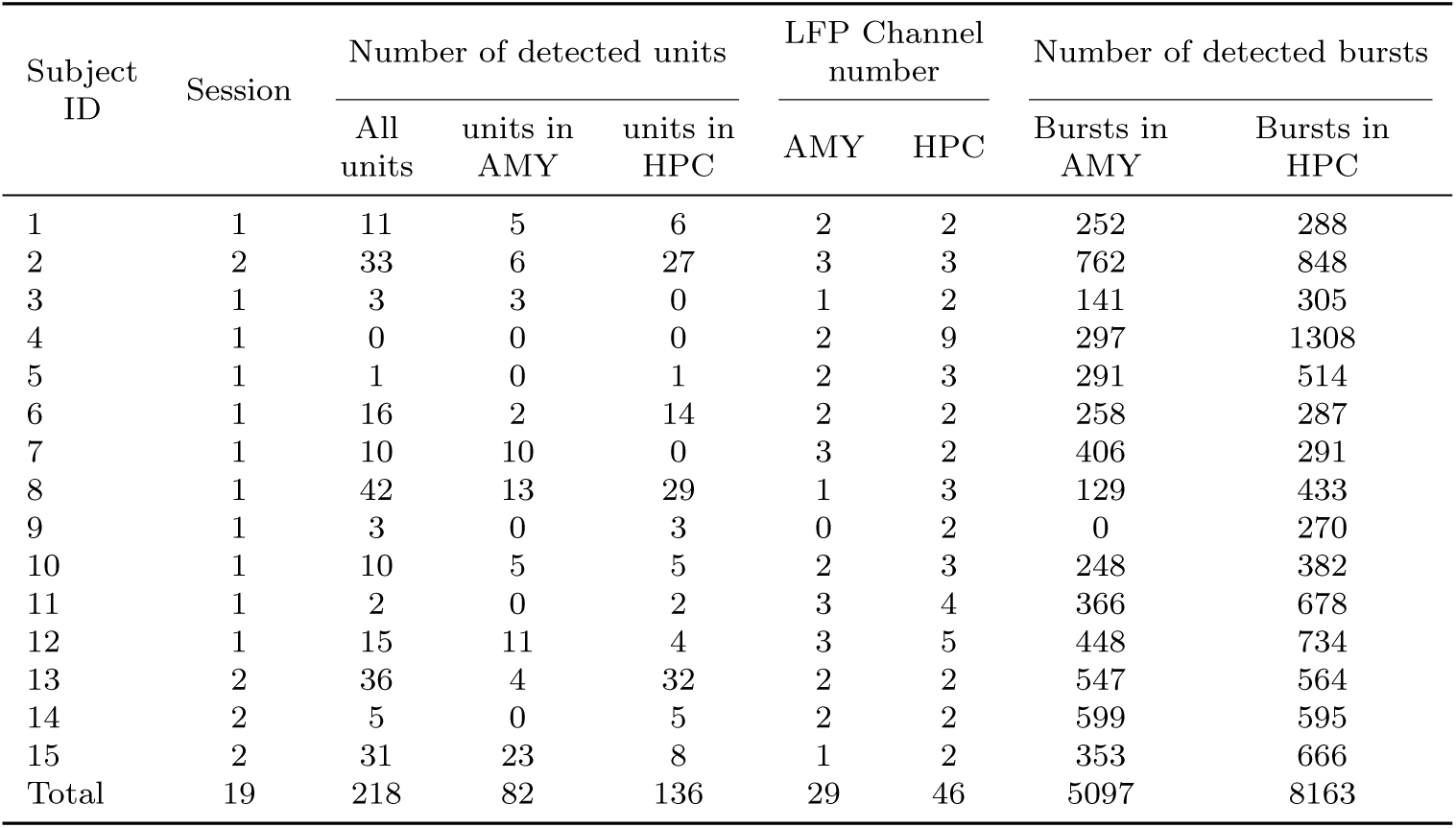
Detected neuron/burst count per region.

| Subject ID | Session | Number of detected units |  |  | LFP Channel number |  | Number of detected bursts |  |
| --- | --- | --- | --- | --- | --- | --- | --- | --- |
|  |  | All units | units in AMY | units in HPC | AMY | HPC | Bursts in AMY | Bursts in HPC |
| 1 | 1 | 11 | 5 | 6 | 2 | 2 | 252 | 288 |
| 2 | 2 | 33 | 6 | 27 | 3 | 3 | 762 | 848 |
| 3 | 1 | 3 | 3 | 0 | 1 | 2 | 141 | 305 |
| 4 | 1 | 0 | 0 | 0 | 2 | 9 | 297 | 1308 |
| 5 | 1 | 1 | 0 | 1 | 2 | 3 | 291 | 514 |
| 6 | 1 | 16 | 2 | 14 | 2 | 2 | 258 | 287 |
| 7 | 1 | 10 | 10 | 0 | 3 | 2 | 406 | 291 |
| 8 | 1 | 42 | 13 | 29 | 1 | 3 | 129 | 433 |
| 9 | 1 | 3 | 0 | 3 | 0 | 2 | 0 | 270 |
| 10 | 1 | 10 | 5 | 5 | 2 | 3 | 248 | 382 |
| 11 | 1 | 2 | 0 | 2 | 3 | 4 | 366 | 678 |
| 12 | 1 | 15 | 11 | 4 | 3 | 5 | 448 | 734 |
| 13 | 2 | 36 | 4 | 32 | 2 | 2 | 547 | 564 |
| 14 | 2 | 5 | 0 | 5 | 2 | 2 | 599 | 595 |
| 15 | 2 | 31 | 23 | 8 | 1 | 2 | 353 | 666 |
| Total | 19 | 218 | 82 | 136 | 29 | 46 | 5097 | 8163 |

